# Interdependent regulation of trabecular meshwork cell physiology and intraocular pressure by KALRN and TMCO1

**DOI:** 10.1101/2025.09.19.677310

**Authors:** Xiaochen Fan, Xu Chen, James W.B. Bainbridge, Maryse Bailly, Ester Reina-Torres, Darryl R. Overby, W. Daniel Stamer, Anthony P. Khawaja, Maria S. Balda, Karl Matter

**Author notes:** Address for Communications: Institute of Ophthalmology, University College London, Bath Street, London EC1V 9EL, UK, /6861, /.

## Abstract

Glaucoma is a leading cause of irreversible blindness. Primary open-angle glaucoma (POAG) is its most common form. Higher intraocular pressure (IOP), resulting from impaired aqueous humour outflow, is the cardinal mediating factor, and all proven treatments aim to lower IOP. POAG is a complex genetic disease with numerous loci linked to POAG and higher IOP. The mechanisms by which risk alleles cause disease remain unclear. Here, using primary trabecular meshwork (TM) cells, a cell type controlling outflow, and mice, we found that the POAG-associated gene *KALRN* encodes an endoplasmic reticulum (ER)-associated Rac regulator essential for TM homeostasis and normal IOP. *KALRN* loss caused widespread disruption from ER to calcium homeostasis and energy metabolism, leading to induction of cell senescence. KALRN-depletion also led to suppression of the ER translocase and calcium regulator TMCO1, encoded by a gene at one of the most significantly associated genomic loci for POAG. TMCO1-depletion *in vitro* and *in vivo* phenocopied KALRN-induced phenotypes, and reduced KALRN expression. These findings establish KALRN and TMCO1 as interdependent regulators of TM homeostasis and IOP, that link regulation of ER and intracellular calcium homeostasis to cell and tissue physiology, and illustrate how different genes linked to glaucoma can form regulatory pathways.

## Introduction

Glaucoma is the leading cause of irreversible blindness globally and primary open-angle glaucoma (POAG) is the most prevalent form of the disease. Higher intraocular pressure (IOP) is the only modifiable risk factor for disease development. Hence, all proven forms of treatment aim to reduce IOP. Increased IOP is caused by reduced aqueous humour and outflow, and is associated with defects in the classical outflow pathway formed by the trabecular meshwork (TM) and Schlemm’s canal (SC). Glaucoma disease development is associated with alterations in TM and SC cells that affect cytoskeletal structure, mitochondrial function, calcium homeostasis, extracellular matrix (ECM) turnover, and stress signalling, as well as SC cell permeability^1–9^. However, how these functions are regulated in outflow pathway cells is not well understood.

Genome-wide association studies (GWAS) have identified hundreds of loci associated with IOP and POAG risk^10–13^. However, little is known about how genes at these loci are functionally linked to IOP regulation and glaucoma. Although TM and SC cells show diverse disease-associated alterations, how these risk genes drive the disease phenotypes or converge on shared regulatory circuits remains largely unknown. Although treatments to lower IOP form the mainstay of management, therapeutic responses vary, and adverse effects are common. Understanding how risk genes function and, particularly, identifying key regulatory mechanisms that control the diverse cellular processes disrupted in glaucoma, would facilitate the development of new, better targeted therapeutic approaches.

Our strategy was to understand how candidate genes at glaucoma-associated loci function by focusing on key signalling mechanisms that govern essential cellular processes. Such a central signalling mechanism is represented by Rho GTPases, which regulate a wide range of cellular processes important for cell structure, morphogenesis, and function^14–17^. Rho GTPases are activated by guanine nucleotide exchange factors (GEFs) that promote GDP/GTP exchange to stimulate Rho GTPase signalling. Perhaps the most intriguing GEF linked to POAG is *KALRN*^11^. *KALRN* encodes a multidomain protein with two GEF domains: one activates Rac1 and RhoG, and the other RhoA^18^. Rac1 and RhoA often oppose each other; hence, a GEF for both Rho GTPases may have multiple cellular functions. KALRN has been mostly studied in neurons, osteoblasts and osteoclasts, in which it was linked to dynamic cellular process requiring actin dynamics^18, 19^.

Here, we investigated the role of KALRN in the regulation of IOP. Using human primary trabecular meshwork cells and a mouse model of KALRN gene depletion, we found that KALRN functions as an endoplasmic reticulum (ER)-associated Rac1 GEF essential for maintenance of normal IOP. Its depletion induced cell-wide changes in TM cells that ranged from disruption of the (ER) and Golgi complex structure to dramatic remodelling of the actomyosin and microtubule cytoskeleton, inhibition of energy metabolism, and induction of cellular senescence. KALRN-depletion also led to disruption of calcium homeostasis and downregulation of *TMCO1*, a gene at a major POAG GWAS locus that is affected by glaucoma-linked tandem repeat polymorphisms^20–23^. TMCO1 is a conserved ER-resident translocase and forms a Ca²⁺ load-activated channel important for maintenance of ER–mitochondrial calcium flux^21, 24^. Depletion of TMCO1 itself led to reduced KALRN levels and phenocopied major loss of function phenotypes of the Rac GEF, including increased IOP and induction of glaucoma-associated histological changes in the mouse TM. These findings identify KALRN and TMCO1 as interdependent key regulators of TM homeostasis and IOP and suggest that a KALRN/TMCO1-based signalling mechanism is essential for ER and calcium homeostasis in TM cells and, thereby, controls cell- and tissue-wide functions including energy metabolism and IOP.

## Results

### KALRN is a functional component of trabecular meshwork cells

Single-cell transcriptomic analyses of outflow-associated tissues suggests that *KALRN* is enriched in the ciliary body and TM cells, with highest expression in the TM cell subtype-beam cell b.(Fig.1A–C)^25^. Immunostaining of non-glaucomatous mouse eyes confirmed strong KALRN expression throughout all TM layers, the inner wall of Schlemm’s canal, and the ciliary epithelium (Fig.1D). Protein–protein interaction mapping (STRING) identified potential functional links between KALRN and proteins encoded by genes at highly significant POAG-associated loci—*TMCO1*, *ABCA1*, *GMDS*, *GAS7*, *TXNRD2*, and *IGF1*^22^ (Fig.1E). Hence, in silico data suggests that *KALRN* may contribute to TM cell physiology and is integrated into glaucoma-associated gene networks.

**Figure 1:**
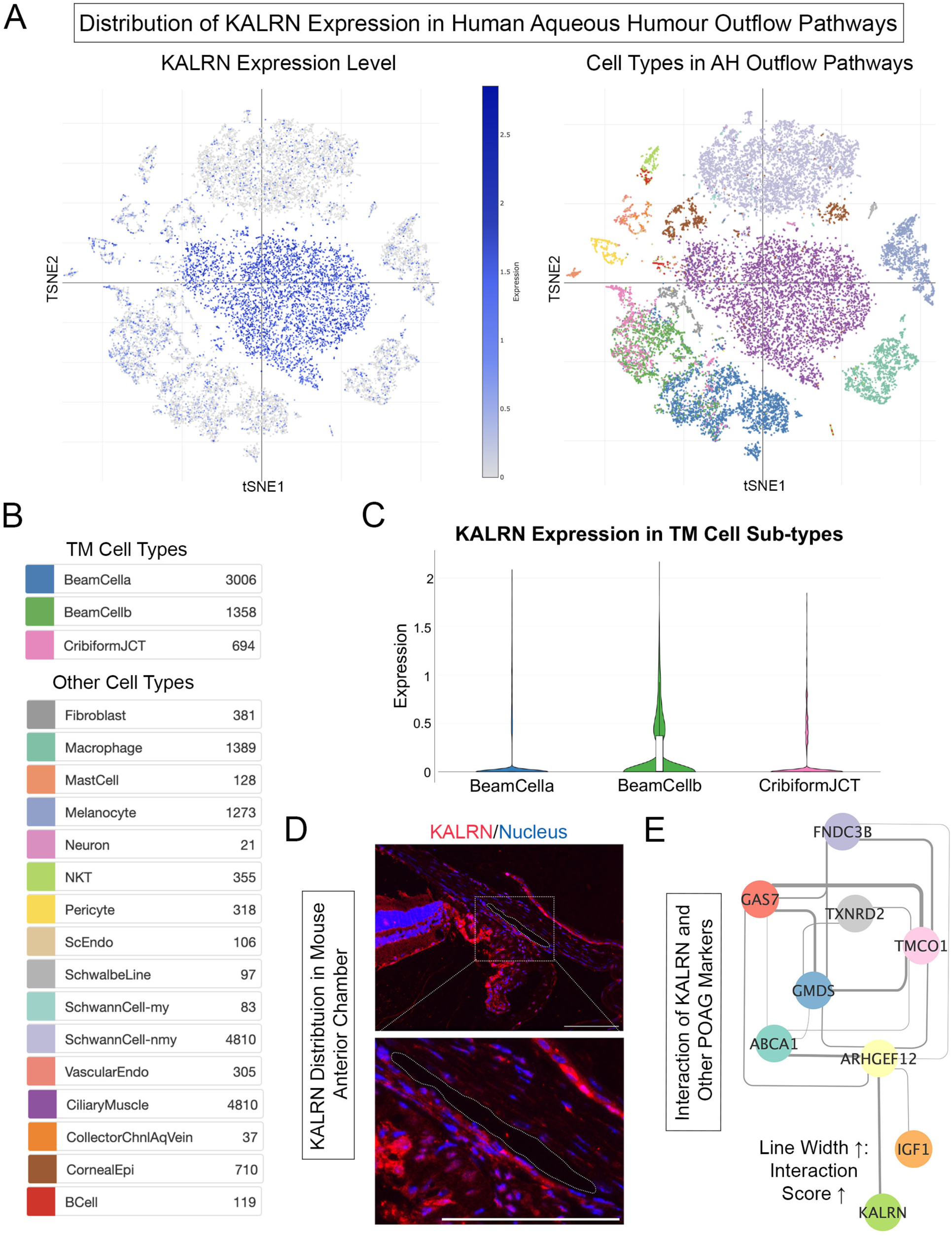
Kalirin localization in aqueous humour outflow tissues and its connection to glaucoma-associated gene networks. **(A–B)** Single-cell RNA sequencing data from human anterior segment tissues revealed KALRN expression enriched in the TM and ciliary body cell populations. Data were adapted from previously published transcriptomic datasets^25^ (https://singlecell.broadinstitute.org/single_cell/study/SCP780/cell-atlas-of-aqueous-humor-outflow-pathways-in-eyes-of-humans-and-four-model-species-provides-insights-into-glaucoma-pathogenesis). t-SNE: t-distributed Stochastic Neighbour Embedding. (C) Within TM subpopulations, KALRN expression is higher in beam cell b than in beam cell a or juxtacanalicular (JCT) cells. **(D)** Immunofluorescence staining of non-glaucomatous adult mouse eyes showed Kalirin protein (red) broadly distributed throughout the TM layers, inner wall of Schlemm’s canal (SC), and non-pigmented ciliary epithelium (NPE). Representative images from 5 eyes. Scale bars, 100µm. **(E)** Protein–protein interaction (PPI) network analysis using STRING illustrates potential functional links between KALRN and established POAG-associated genes (TMCO1, ABCA1, GMDS, GAS7, TXNRD2, and IGF1). Increased line width indicates the higher interaction score from String.

We setup a loss of function approach to determine the importance of KALRN in TM cells. Three siRNAs targeting *KALRN* sequences within different domains were synthesized (Supplementary Fig.1A). The three *KALRN* siRNAs were then transfected individually or as a pool into primary human TM cells. A non-targeting siRNA was used as a control. All *KALRN* siRNAs significantly reduced KALRN-12 protein levels, the main isoform we could detect in TM cells, as measured by immunoblotting (Supplementary Fig.1B-C). In human TM cells, all three siRNAs transfected alone or as a pool induced consistent morphological alterations, resulting in a loss of the typical extended cell morphology to a more spread appearance (Supplementary Fig.1D-E). Thus, *KALRN* expression can be effectively reduced, and the protein is important for the maintenance of TM cell morphology.

Further analysis of KALRN-depleted cells revealed that the depletion induced striking structural and functional changes in four different strains of primary human TM cells. Control siRNA transfected TM cells exhibited little cell-cell adhesion, similar to non-transfected cells; in contrast, KALRN-depleted cells formed adherens junction-like structures containing components such as ZO-1, α-catenin, N-cadherin, and I-Afadin (Fig.2A-C). More than 80% of KALRN-depleted cells exhibited adherens junction-like structures with continuous ZO-1 staining along more than half of the cell’s long axis (Fig.2B). KALRN-depleted cells spread more and became more rounded in shape (Fig.2D). KALRN-depletion leads to a strong reduction in expression of the proliferation marker Ki67 (Fig.2E-F), and to significantly larger nuclei, which may suggest cell cycle arrest or senescence (Fig.2G). Cell spreading and increased cell adhesion is often associated with cell senescence. Consistent with this, senescence associated-β-galactosidase (SABG) staining revealed strong increases in the proportion of SABG–positive cells in KALRN-depleted TM cells, indicating increased cell senescence in all four analysed TM cell strains (Fig.2H-I). These features indicate that KALRN maintains TM cell morphology and prevents cell senescence.

**Figure 2:**
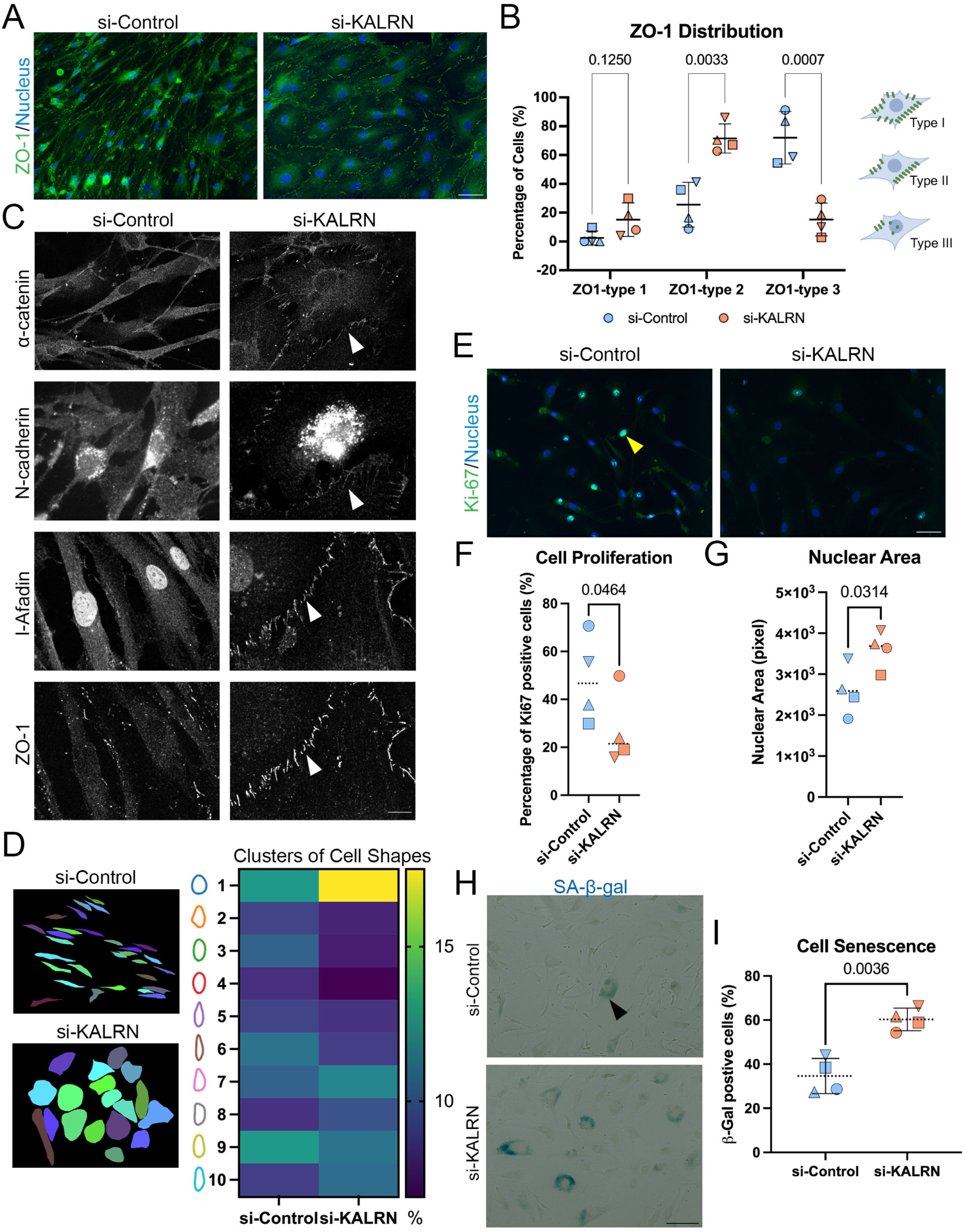
KALRN depletion induces remodelling of cell junctions and induces senescence in human TM cells. **(A)** Immunofluorescence staining of ZO-1 (green) in TM cells following KALRN knockdown showed altered junctional organization (n=4 donors). Scale bar, 50µm. **(B)** KALRN knockdown led to an increase in ZO-1 Type-2 junctions, defined by continuous staining along >50% of the cell’s long axis. Quantification showed elevated frequencies of both Type-2 and Type-3 (continuous staining along <50% of the cell’s long axis) ZO-1 patterns relative to controls (n=4 donors; paired t-tests). **(C)** Co-localization of ZO-1 with adherens junction proteins α-catenin, N-cadherin, and I-Afadin (grey) at cell boundaries (white arrows) in KALRN-depleted cells (n=2 donors). Scale bar, 50µm. **(D)** Shape segmentation of TM cells based on principal component analysis (PCA) revealed 10 distinct morphological clusters. Heatmap represents the relative frequency of cell shapes in control and KALRN-depleted populations (brighter colour indicates higher proportion) (si-Control n=737 cells and si-KALRN n=480 cells from 4 donors). **(E-F)** KALRN depletion reduced cell proliferation, as determined by percentage of Ki-67 (green) positive cell (yellow arrow) counting (n=4 donors; paired t-test). Scale bar, 50µm. **(G)** Nuclear area was enlarged in KALRN-depleted cells (n=4 donors; paired t-test). **(H-I)** Senescence-associated β-galactosidase (SAβG) staining (blue) showed increased β-gal– positive cells (black arrow) in the KALRN knockdown group (n=4 donors; paired t-test). Scale bar, 50µm.

### KALRN regulates the actin and microtubule cytoskeleton

Depletion of KALRN induces increased cell-cell adhesion and cell spreading, processes regulated by the cytoskeleton (Fig.2). KALRN is a GEF for the Rho GTPases Rac and RhoA, major regulators of the cytoskeleton and cell-cell adhesion^26, 27^. Hence, we next asked if KALRN impacts on the cytoskeleton.

In control TM cells, F-actin was found to assemble fibres that were decorated by active myosin as indicated by staining with antibodies against mono-(p-MLC2, Ser19) and di-phosphorylated (pp-MLC2, Thr18/Ser19) myosin light chain 2. In KALRN-depleted cells from four donors, both p-MLC2 and pp-MLC2 levels were reduced, as shown by immunostaining and western blotting (Fig.3A-C, Supplementary Fig.2A). Hence, actomyosin contractility might be impaired by KALRN-depletion.

**Figure 3:**
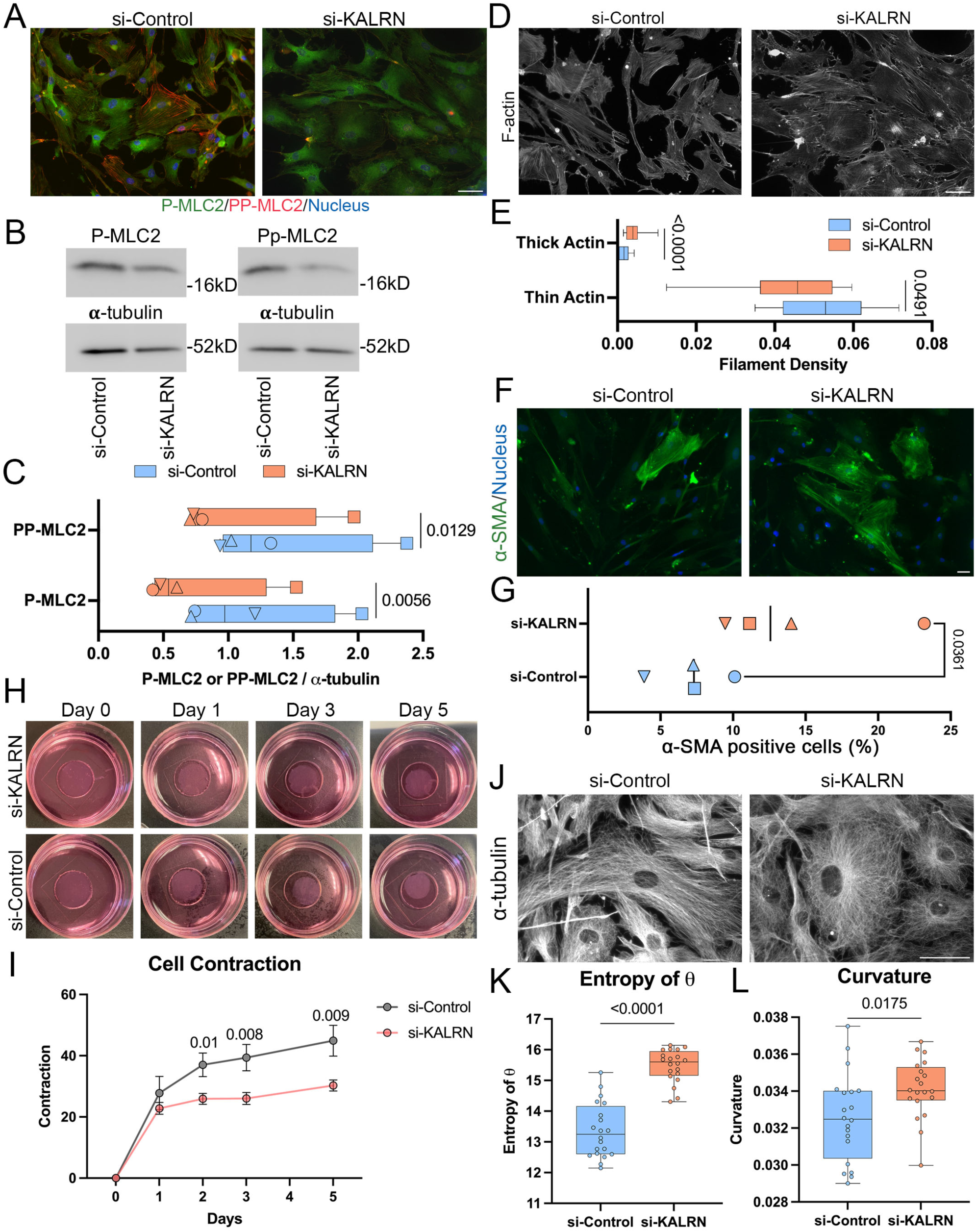
KALRN depletion induces cytoskeletal reorganization and alters cell contractility. **(A)** Immunofluorescence staining of phosphorylated MLC2 (p-MLC2, green) and diphosphorylated MLC2 (pp-MLC2, red) showed reduced signal intensity in KALRN-depleted cells; n=4 donors, Scale bar, 50µm. **(B-C)** Western blot analysis confirms decreased p-MLC2 (MW: ∼20kDa) and pp-MLC2 (MW: ∼20kDa) expression following KALRN depletion (WB membrane: TM219; paired t-tests; n=4 donors). **(D-E)** Phalloidin staining (grey) showed increased thick filament density and decreased thin filament density after KALRN depletion (n=20 cells per group, TM155; Mann–Whitney test). Scale bar, 50µm. **(F-G)** Immunofluorescence reveals increased numbers of α-SMA–positive cells (green) in KALRN-depleted cultures (n=4 donors; paired t-test). Scale bar, 50µm. **(H-I)** Collagen gel contraction assays indicate reduced contractility in KALRN-depleted cells from day 1 to day 5 (n=3 replicates of TM219). Consistently replicated in TM155 and TM213. **(J)** α-Tubulin immunostaining (grey) reveals disorganization of microtubule networks in KALRN-depleted cells (n=4 donors). Scale bar, 50µm. **(K)** Microtubule disorganization was quantified by calculating the angular entropy of α-tubulin filament orientation relative to the radial axis (8). KALRN-depleted cells induced higher angular entropy, indicating increased network disorganization (n=20 cells per group, TM155; Welch’s t-test). **(L)** Microtubule curvature was increased in KALRN-depleted cells (n=20 cells per group, TM155; unpaired t-test).

Phalloidin staining of F-actin in KALRN-depleted cells showed cross-linked actin network (CLAN)-like structures, a cytoskeletal phenotype previously reported to be induced in TM cells in glaucoma models^7^. (Supplementary Fig.2B). Quantitative analysis of phalloidin-stained F-actin fibres, segmented by width into thin (1–2 pixels) and thick (3–15 pixels) categories (Supplementary Fig.2C), revealed a significant reduction in thin fibres and a corresponding increase in thick fibre density in KALRN-depleted cells (Fig.3D-E). Thickening of F-actin fibres often occurs in cells that become more contractile. Increased contractility during TM to mesenchymal transition in glaucoma is associated with increased expression of α-smooth muscle actin (α-SMA). Upon KALRN-depletion, the percentage of α-SMA positive cells indeed increased, indicating that the cells became more mesenchymal (Fig.3F-G).

The decreased MLC2 phosphorylation suggested that the cells became less contractile; however, the changes in F-actin organisation and α-SMA expression implied increased contractility. Therefore, we measured overall cellular contractility using a collagen gel contraction assay^28^. Figure 4H-I shows that KALRN-depletion led to a decrease in contractility. Interestingly, the reduction in contractility was more pronounced in TM cells derived from an older donor compared to those from younger donors (Supplementary Fig.2D), suggesting age-dependent sensitivity to KALRN loss.

**Figure 4:**
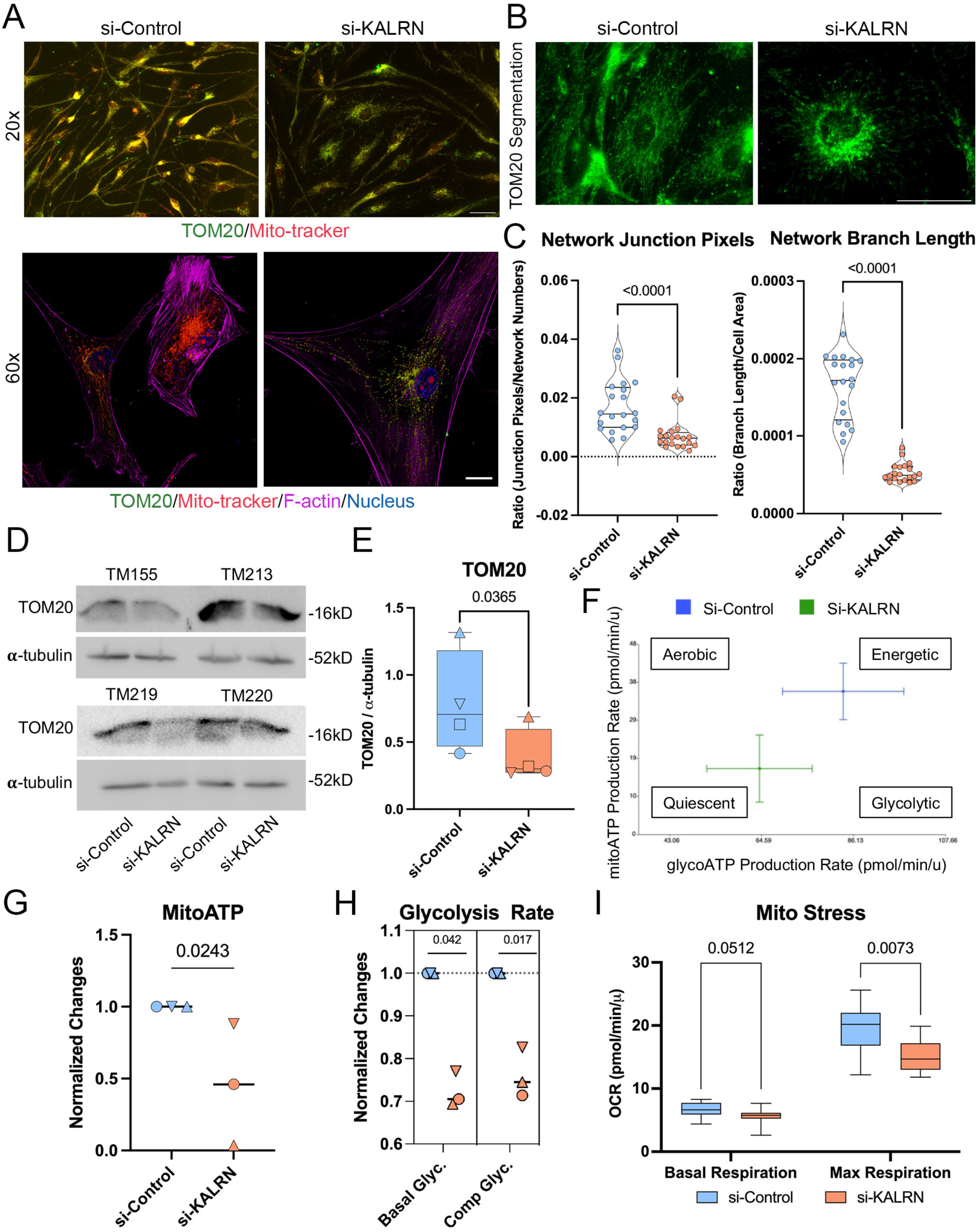
KALRN depletion disrupts mitochondrial structure and impairs cellular energy metabolism. **(A)** Immunostaining for TOM20 (green) and Mito-tracker (red) showed fragmented and disorganized mitochondrial networks in KALRN-depleted TM cells (n=4 donors). Purple: F-actin stained by phalloidin. Scale bars, 50μm and 20μm; **(B-C)** Quantification of mitochondrial network structure stained with TOM20 (green) revealed reduced junction pixel-to-network ratios (fold change relative to control) and decreased branch length normalized to cell area (n=20 cells per group; Mann–Whitney test). **(D–E)** Western blot and quantification confirm reduced TOM20 protein (MW:16kDa) levels in KALRN-deficient cells (n=4 donors: paired t-test). **(F)** Seahorse assay–derived energy maps showed reduced overall energetic and glycolytic capacity in KALRN-depleted cells (green) compared with controls (blue); TM155. Consistently replicated in TM213 and TM219. **(G)** Seahorse assay showed a reduction in mitochondrial ATP production (n=3 donors: TM155, TM213 and TM219; paired t-test). **(H)** Seahorse assays revealed basal and compensatory glycolysis rates are reduced in KALRN-deficient cells (n=3 donors; paired t-test). **(I)** Basal and maximum mitochondrial respiration are reduced in KALRN-deficient cells (TM155; n=12; Mann–Whitney test).

Similar to the actin cytoskeleton, the microtubule network regulates cell shape changes and is anchored to cell-cell adhesions^29^. Depletion of KALRN led to major changes in microtubule organization. In control TM cells, microtubules formed parallel arrays along the long axis of the cells (Fig.3J). Upon KALRN-depletion, the microtubule networks adapted a structure more typical for mesenchymal cells, radiating from an apparent perinuclear microtubule organising centre towards the cell periphery, often apparently bending or overlapping. To measure this change in microtubule architecture, we calculated the entropy of the angle (θ) between the direction of each microtubule segment and the cell’s radial axis. Cells lacking KALRN showed a significant increase in angular entropy, indicating a loss of directional alignment among microtubules (Fig.3K), as well as increased curvature (Fig.3L; see Supplementary Fig.2E-G for segmentation examples and schematic diagrams explaining the calculation of θ and entropy). These findings highlight KALRN’s role in maintaining the overall cytoskeletal organization characteristic of TM cells, which is critical determinant of TM contractility and flow regulation.

We next asked whether these cytoskeletal changes have functional consequences. We first measured cell migration using live-cell imaging over two hours, quantifying migration distance and cell shape changes. The primary TM cells migrated only slowly, and migration was further reduced upon KALRN-depletion (Supplementary Fig.3A-C, supplementary videos 1 and 2). Similarly, measurements of Hausdorff distances — a measure of maximal deviation between initial and final cell outlines — revealed significantly less morphological changes in migrating KALRN-depleted cells (Supplementary Fig.3D-F), indicating impaired dynamic shape remodelling.

KALRN is thus required for normal structure and organisation of the actin and microtubule cytoskeleton, as well as dynamic cellular processes requiring cytoskeletal remodelling and dynamics.

### KALRN regulates energy metabolism

The cytoskeletal phenotype of KALRN-depletion displays an apparent discrepancy between induction of a cytoskeletal architecture commonly associated with increased contractility (i.e., thicker F-actin bundles and increased α-SMA expression) but decreased MLC2 phosphorylation and reduced contractility. Hence, the reduced contractility might be caused by downregulated metabolic activity.

To assess mitochondrial integrity of KALRN-deficient cells, we first performed immunostaining for TOM20, an outer mitochondrial membrane protein^30^, and MitoTracker, a membrane potential–sensitive fluorescent dye that labels active mitochondria^31^. TOM20 staining revealed significant disruption and fragmentation of the mitochondrial network, evidenced by reduced junction pixel-to-network ratios and shorter branch lengths normalized to cell area (Fig.4A-C; see Supplementary Fig.4A for the segmentation of TOM20-labelled mitochondrial networks and Supplementary Fig.4B for schematics of the quantification method). Western blot analysis showed reduced TOM20 protein levels, suggesting decreased mitochondrial mass after KALRN-depletion (Fig.4D-E). The protein level of dynamin-related protein 1 (DRP1), a mitochondrial fission marker, was slightly increased (Supplementary Fig.4C–D), suggesting enhanced mitochondrial fission activity.

The MitoTracker results suggest that KALRN-depletion inhibits mitochondrial metabolism. Therefore, we next employed Seahorse assays to measure oxidative phosphorylation and glycolysis in TM cells upon depletion of KALRN. Measurements of ATP production indicated that mitochondrial ATP production were downregulated by KALRN-depletion (Fig.4F-G). Assays to measure maximum capacity of glycolysis and oxidative phosphorylation further revealed that both basal and maximum rates of both metabolic pathways were downregulated; however, there was considerable variation between the different cell strains (Fig.4H-I; Supplementary Fig.4E–F). Additionally, mitochondrial proton leak significantly increased in cells from an old donor but not in the cells from young donors (Supplementary Fig.4G), indicating that remaining mitochondria were functionally compromised. Intriguingly, maximal respiration decreased in young donor cells, which maintained a low proton leak. In old donor cells, however, maximal respiration was unaffected, but the proton leak increased by a factor of ∼3. Hence, young and old cells had decreased mitochondrial ATP production capacities by KALRN-depletion. Despite this, mitochondrial activation—measured as the ratio of MitoTracker to TOM20 staining^32^ — was unchanged (Supplementary Fig.4H), indicating that reduced oxidative phosphorylation mainly resulted from decreased mitochondrial mass, as indicated by the reduced TOM20 expression, rather than loss of membrane potential in KALRN depleted cells.

### KALRN-depletion disrupts autophagy

The low ATP production rates of both glycolysis and oxidative phosphorylation should lead to increased AMP levels and, hence, activation of AMP-activated protein kinase (AMPK) and, in turn, activation of autophagy to simulate metabolism^33^. Indeed, phosphorylation of AMPK increased in all cell strains analysed but was more pronounced in younger cells with low AMPK phosphorylation levels under control conditions (Fig.5A-B). As metabolism was low in TM cell strains from old and young donors upon KALRN-depletion, we asked whether autophagy is regulated by KALRN.

**Figure 5:**
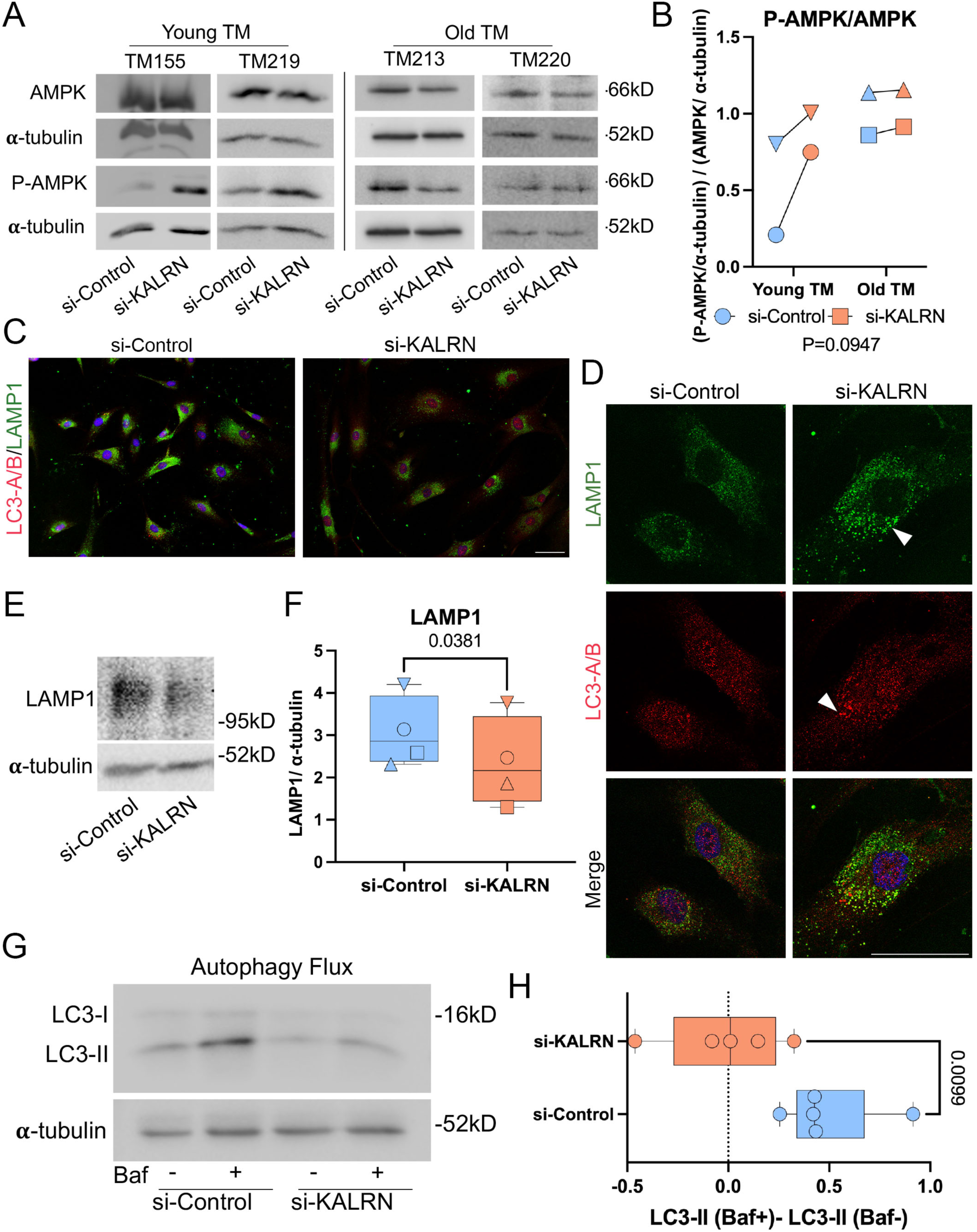
KALRN depletion disrupts autophagic flux and leads to intracellular accumulation of fibronectin. **(A–B)** Western blot analysis of AMPK and phosphorylated AMPK (p-AMPK, ∼62kDa) in control and KALRN-depleted TM cells. Blots and quantification showed increased p-AMPK/AMPK ratios following KALRN depletion, with a greater effect in young donors compared to aged donors. Data from four donor strains are shown ([●, ▼], young; [▲, ∎], aged). Paired t-test. **(C–D)** Immunofluorescence staining of LC3 (autophagosome marker, red) and LAMP1 (lysosomal marker, green) reveals enlarged puncta (white arrow) in KALRN-depleted TM cells (n=3 donors; TM155, TM213, and TM219). Scale bars, 50μm. **(E–F)** Western blot analysis showed reduced LAMP1 protein levels in KALRN-deficient cells (n=4 donors; Paired t-test). **(G–H)** Treatment with Bafilomycin A1 results in decreased LC3-II protein (∼14kDa) accumulation in KALRN-depleted cells compared to controls (n=5 technique replicates, TM155; Paired t-test), indicating impaired autophagic flux.

We first immunolabelled LC3, an autophagosome marker, and LAMP1, a major lysosomal transmembrane protein known to regulate autophagy^34, 35^. KALRN-deficient TM cells exhibited enlarged LC3- and LAMP1-positive puncta (Fig.5C-D), and reduced LAMP1 protein levels by immunoblotting (Fig.5E-F), suggesting that autophagy may be affected by KALRN-depletion. Assessments of autophagic flux based on measurements of LC3 levels upon Bafilomycin A1-mediated inhibition of LC3 degradation^36^ revealed diminished LC3 accumulation in *KALRN*-depleted cells compared to controls (Fig.5G-H). Thus, KALRN-depletion disrupts autophagy.

### KALRN is required for endoplasmic reticulum and calcium homeostasis

Immunofluorescence of KALRN revealed a reticular distribution similar to that of ER markers such as Calnexin^37^ and CLIMP-63^38^, suggesting association with the ER (Fig.6A). Upon KALRN knockdown, ER morphology was markedly disrupted, with vacuole-like expansions observed by staining for CLIMP-63 and TMCO1^21^ (Fig.6A). BiP, a major ER-resident chaperone involved in protein folding^39^, showed decreased expression and formed punctate structures (Fig.6A). Reduced BiP and TMCO1 levels upon KALRN-depletion in all four TM cell strains were confirmed by immunoblotting (Fig.6B-C, Supplementary Fig.5A). *TMCO1* is a major glaucoma-associated gene, and *TMCO1* variants have been linked to increased IOP in POAG^20, 22, 23^. Hence, KALRN and TMCO1may be functionally linked.

**Figure 6:**
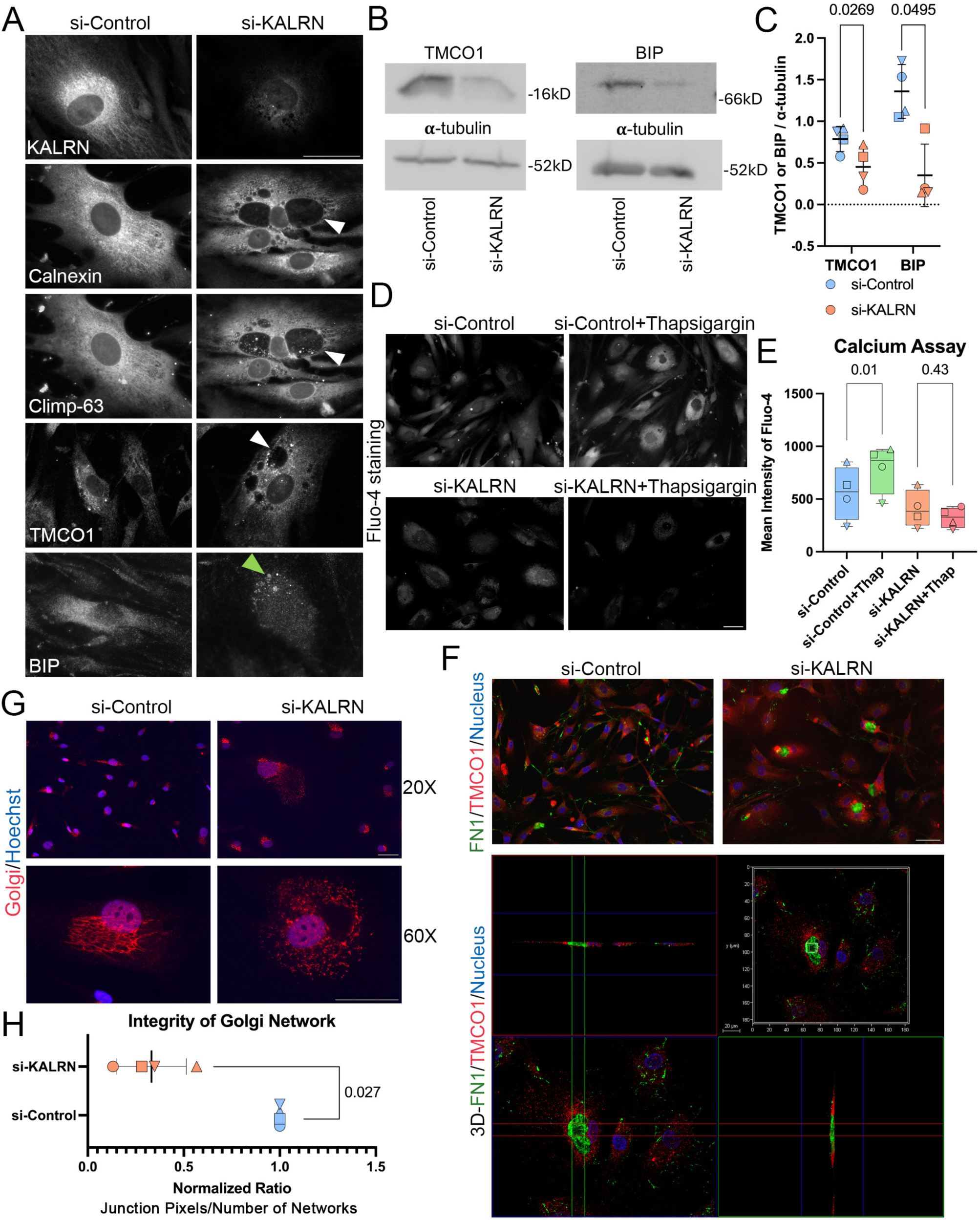
KALRN is required for ER morphology and function and impairs calcium homeostasis. **(A)** Immunofluorescence staining showed Kalirin exhibited a reticular distribution. Vacuole-like expansions were observed in the KALRN depleted cells stained with Calnexin, CLIMP-63, TMCO1, and KALRN (grey). BiP (grey) displayed reduced staining intensity and punctate localization. White arrows: vacuole-like expansions. Green arrows: BIP puncta. n=4 donors; Scale bar, 50µm. **(B-C)** Western blot analyses showed reduced BiP and TMCO1 protein levels in KALRN-depleted cells (n=4 donors: young donors; [▲, ∎]: old donors; paired t-tests. **(D-E)** ER calcium release was assessed using Thapsigargin. In control cells, Thapsigargin induced Fluo-4-stained cytosolic calcium release. In KALRN-depleted cells, the Fluo-4-stained calcium decreased. Scale bar, 50µm; n=4 donors; paired t-test. **(F)** Immunofluorescence showing FN1 (green) in vacuole-like expansions of TMCO1-stained TM cells (red) with KALRN depletion. TM219; Top: FN1 and TMCO1 staining; scale bar, 50µm; Bottom: 3D-reconstructed FN1 and TMCO1 staining; scale bar, 20µm. **(G)** Immunofluorescence showing disrupted Golgi structure in KALRN depleted TM cells stained with GOLPH2/GP73 (red); n=4 donors; scale bar, 50µm. **(H)** Quantification of Golgi structure integrity revealed reduction in network integrity in KALRN depleted cells (n=4 donors; paired t-tests for not normalized data).

The observed morphological disruption of the ER and the suppression of genes important for calcium homeostasis and components required for membrane insertion and protein folding, suggests that KALRN-depletion has a major impact on ER function. Given the impact on TMCO1, we first measured calcium release upon Thapsigargin treatment. Figure 6D-E shows that KALRN-depletion abolished Thapsigargin-induced Ca²⁺ release, suggesting empty ER calcium stores. These results indicate that KALRN is required for calcium homeostasis.

Several of the ER markers found to be disrupted in KALRN-depleted cells are required for membrane integration into or protein folding in the ER (e.g., TMCO1, BiP, calnexin). Hence, synthesis and transport of membrane and luminal/secreted proteins may be affected. Fibronectin 1 (FN1), a major extracellular matrix protein secreted by TM cells, was found to accumulate in the vacuoles induced upon KALRN-depletion (Fig.6F, Supplementary Fig.5B-C), supporting a defect in ER protein export. Similarly, some vacuoles detected by labelling calnexin had luminal staining, indicating accumulation of the soluble isoform^40^ (Supplementary Fig.5D). Disruption of ER function can lead to disintegration of the Golgi complex^41^. Depletion of KALRN indeed disrupted the Golgi network (Fig.6G-H). Hence, KALRN is required for maintenance of both the ER and the Golgi complex.

The association of KALRN with the ER and the disruption of the ER upon KALRN-depletion suggests that ER malfunction contributes to many of the observed KALRN-depletion phenotypes. Apart of the ER’s role in protein secretion and secretory pathway maintenance, the ER contributes lipids for autophagosome formation, and ER calcium stores are important to maintain normal energy metabolism.

### KALRN signals via Rac in TM cells

KALRN has different structural domains with specific functions (Supplemental Fig.1A)^18, 42^. Two of these domains have guanine nucleotide exchange factor activity: Rho-GEF1 and Rho-GEF2. Rho-GEF1 has been linked to Rac1 and RhoG signalling whereas RhoGEF2 can activate RhoA.

To determine whether KALRN re-expression could rescue the cellular phenotypes induced by its loss, KALRN-depleted human TM cells were transfected with rat c-Myc–tagged KALRN-7, a short isoform containing only one Rho-GEF domain, or KALRN-12, the main isoform expressed in TM cells (Supplemental Fig.1A). Immunostaining confirmed successful expression of both isoforms in transfected cells (Fig.7A). Expression of either KALRN-7 or KALRN-12 led to a reduction in cell size and reduced ZO-1 localization at cell–cell junctions (Fig.7B-D). KALRN re-expression thus rescued main phenotypes induced by KALRN knockdown. KALRN-7 did so very effectively, indicating that domains C-terminal to the Rho-GEF1 domain are not required for the analysed functions in TM cells and, hence, excludes domains such as the RhoA-specific Rho-GEF2 and the kinase domain.

**Figure 7:**
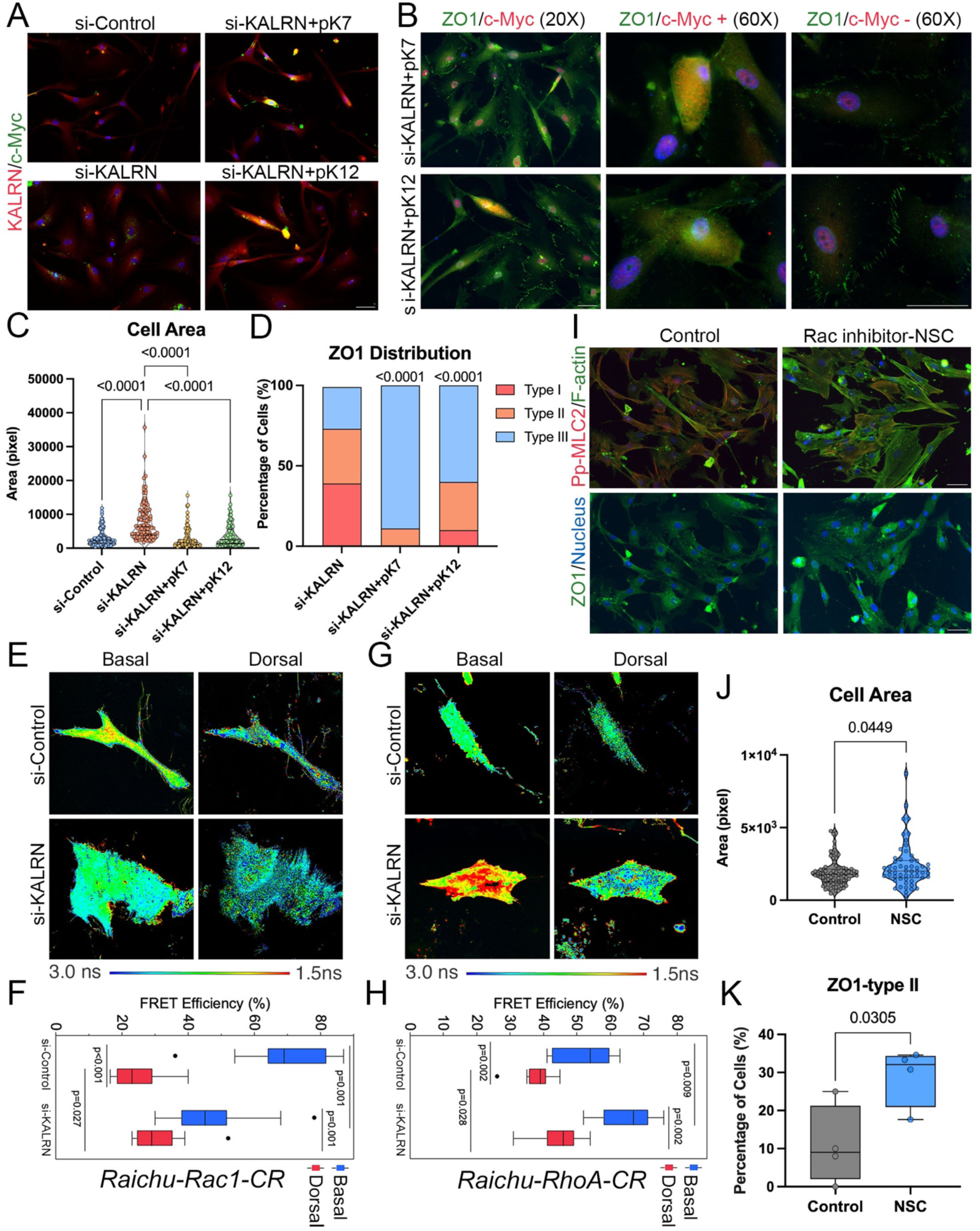
KALRN regulates while Rac1 activation. **(A)** KALRN-depleted TM cells transfected with c-Myc–tagged KALRN-7 or KALRN-12 (TM219) showed successful isoform expression (green: c-Myc, red: KALRN). Scale bar, 50μm. **(B)** ZO-1 immunostaining (green) decreased at cell-cell junctions in KALRN-7 and KALRN-12 restored cells (c-Myc–positive cells, red). Scale bars, 50μm. **(C)** Quantification confirmed reduction in cell area after re-expression of KALRN-7 and −12 isoforms (n=30-50 cells per group; Kruskal–Wallis test; Mean areas: si-Control: 1868 pixel², si-KALRN: 7242 pixel², si-KALRN + plasmid KALRN-7 (pK7): 3716 pixel², si-KALRN + plasmid KALRN-12 (pK12): 4603 pixel²). **(D)** Chi-square analysis showed the changed junctional ZO-1 patterns in the cells with re-expression of KALRN-7 and −12 isoforms. TM219; si-KALRN=76 cells; si-KALRN + pK7=18 cells; si-KALRN + pK12=10 cells. **(E-H)** FLIM–FRET analysis showed reduced Rac1 activity and increased RhoA activation in KALRN-depleted cells (TM219). In Rac1 biosensor–expressing cells, warmer colours (shorter lifetimes) indicate higher activity. KALRN depletion reduced Rac1 activity in the basal half of the cells, as shown by cooler colour shifts. RhoA activity increased at the basal side, indicated by warmer colours (Kruskal-Wallis tests; Rac1, n=13 per group; RhoA, n=10 per group). **(I)** Inhibition of Rac using NSC23766 in TM cells reproduced features of KALRN loss, including decreased pp-MLC2 staining (red), thicker F-actin filaments (green), and increased ZO-1 staining at cell-cell junction area. TM219. Scale bar, 50μm. **(J)** Rac inhibition increased cell area (TM219; control: n=76 cells, NSC: n=55 cells from 5 images; Mann–Whitney test). **(K)** Percentage of TM cells with ZO-1 type II distribution were elevated following Rac inhibition (n=4 images, TM219; unpaired t-test).

The Rho-GEF1 domain can activate Rac1signalling. We first asked whether Rac1 activity is affected by the depletion of KALRN using a FRET-based biosensor for Rac1. We also monitored RhoA activity using an analogous sensor. Both sensors are based on the original Raichu sensor constructs in which the fluorescent FRET pair was replaced by Clover and mRuby2 to increase the dynamic range^43, 44^. Figure 7E-H shows that most Rac1 activity was detected in the basal half of the cells using FLIM to assess the FRET efficiency. Depletion of KALRN led to a reduction in the FLIM-FRET efficiency by more than a third, indicating decreased Rac1 activity. In contrast, RhoA activity increased, likely contributing to the cell spreading observed after KALRN-depletion. KALRN is thus required for normal Rac1 activation.

To investigate whether Rac1 inhibition causes effects induced by KALRN-depletion, we treated TM cells with the Rac1-specific inhibitor NSC23766^45^. Rac1 inhibition resulted in reduced pp-MLC2, thicker actin stress fibres, enlarged cell size, enhanced ZO-1 localization at cell-cell junctions, formation of ER vacuoles, and Golgi structure disruption (Fig.7I-K, Supplementary Fig.6A-C). Hence, inhibition of Rac1 replicates phenotypes induced by KALRN-depletion.

Our data thus indicate that Rac1 is a major effector of KALRN in TM cells, and that disruption of Rac1 signalling contributes to the KALRN-depletion-induced phenotype.

### In vivo depletion of KALRN increases intraocular pressure and induces glaucoma-associated TM cell phenotypes

We next aimed to test the role of KALRN in the TM *in vivo*. Mouse *KALRN* targeting Accell siRNAs, which are taken up by cells without transfection reagents, were first tested *in vitro* for depletion of KALRN protein by immunoblotting (Supplementary Fig.7A). Each animal was then injected with control Accell siRNA in one eye and *KALRN*-targeting siRNA in the other eye (alternating left and right). Immunohistochemistry revealed that KALRN expression was downregulated in the TM between days 4 to 8 and then recovered (Fig.8A-B, Supplementary Fig.7B). Thus, KALRN expression could be successfully reduced *in vivo*.

**Figure 8:**
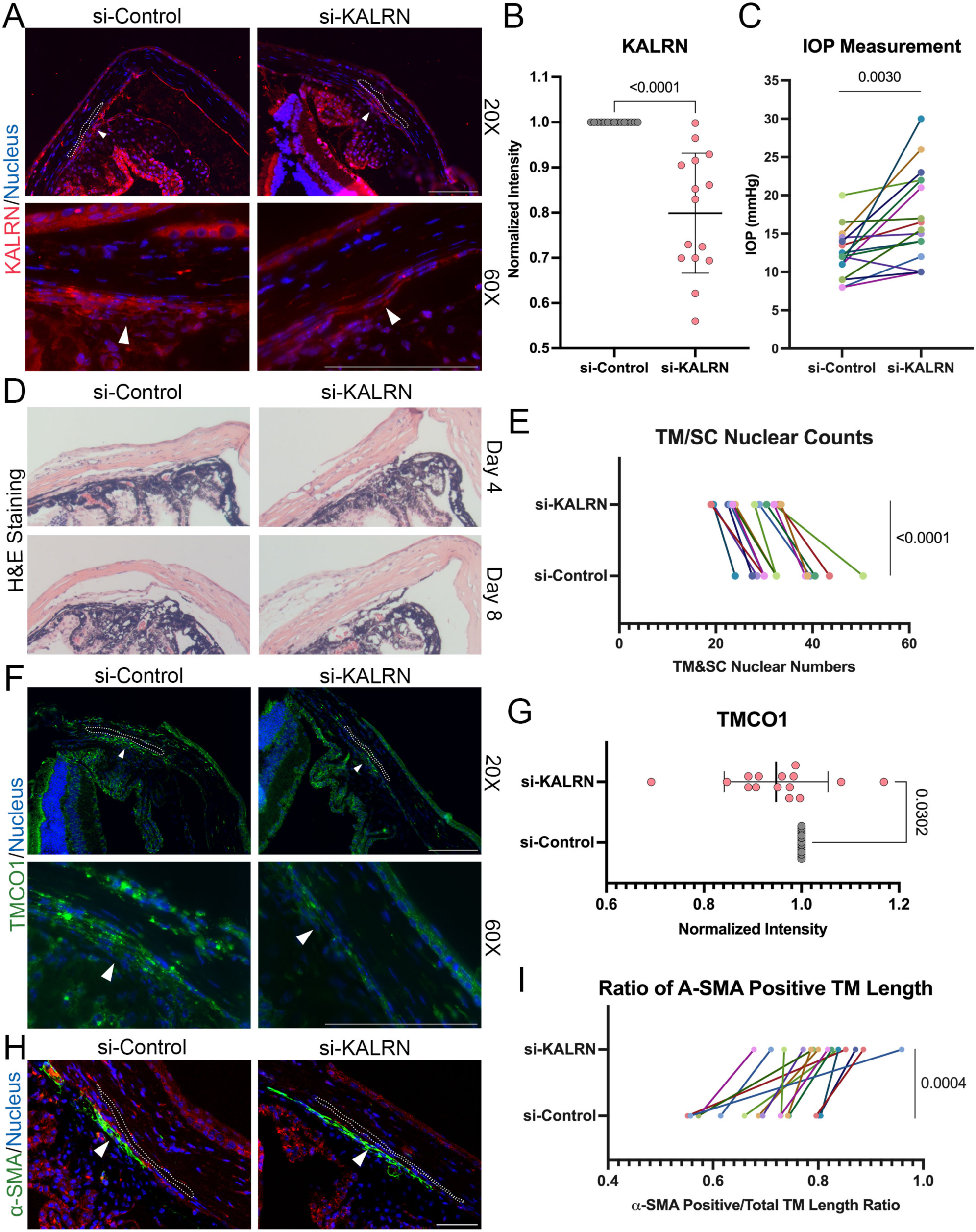
KALRN depletion increases intraocular pressure and induces cellular loss in the TM of mouse eyes. **(A-B)** Immunostaining confirmed reduced KALRN expression (red) in the TM tissue in mice with KALRN siRNA injection (n=15; Wilcoxon matched-pairs signed rank test). White arrow: TM tissue; dashed circle outlines Schlemm’s canal (SC). Scale bars, 100μm. **(C)** IOP was elevated in mouse eyes injected with KALRN siRNA compared to control siRNA injected eyes (n=15 mice; paired t-test). **(D–E)** H&E staining showed structural alterations in KALRN-depleted eyes, including loss of TM cells and increased TM pigmentation. Cell density was reduced (n=15 mice; paired t-test). Scale bar, 50μm. **(F-G)** TMCO1 expression (green) was reduced in TM tissue following KALRN depletion, consistent with in vitro findings (n=15 mice; Wilcoxon matched-pairs signed rank test); White arrow: TM tissue; Dashed circle outlines SC. Scale bar, 100μm. **(H–I)** α-SMA immunostaining (green) showed broader distribution in KALRN-depleted TM, extending from the uveal meshwork toward the corneal endothelium transition zone. White arrow: TM tissue; dashed circle outlines SC. Scale bar, 50μm. Quantification of α-SMA–positive TM length revealed an increase (n=15 mice; paired t-test).

IOP measurements by rebound tonometry revealed that KALRN knockdown caused a significant increase in IOP compared to control siRNA injections into the contralateral eye (Fig.8C). Histological analysis using haematoxylin and eosin (H&E) staining revealed structural changes in the TM, including fewer TM and Schlemm’s canal inner wall cells and increased pigmentation (Fig.8D-E). TMCO1 levels were reduced in KALRN-depleted TM tissue, consistent with the *in vitro* data (Fig.8F-G).

α-SMA staining was mainly confined to the inner TM layer near the iris angle (uveal meshwork) in control eyes. Following KALRN-depletion, α-SMA–positive cells extended from the iris angle toward the transition zone between the TM and the corneal endothelium, showing a broader α-SMA distribution within the TM with KALRN-depletion (Fig.8H-I; see Supplementary Fig.7C for explanation of the quantification). Correlation analysis further revealed that α-SMA staining was associated with pigmentation levels (Pearson’s coefficient; Supplementary Fig.7D-F).

These findings indicate that KALRN loss in vivo increases IOP and disrupts TM homeostasis, inducing phenotypes that we also observed *in vitro* and that are associated with high pressure glaucoma.

### TMCO1 suppression mimics KALRN-depletion phenotypes

TMCO1 is a major glaucoma-associated gene linked to elevated IOP^20, 22, 23^. TMCO1 is downregulated by KALRN-depletion and is important for calcium homeostasis. Hence, we asked whether reduced expression of TMCO1 contributes to the KALRN phenotype.

Efficient TMCO1 knockdown was achieved using distinct siRNAs in primary human TM cells and a pool of Accell siRNA in mouse cells (Supplementary Fig.8A-B). In primary human TM cells, TMCO1 knockdown induced cellular changes similar to those observed upon KALRN-depletion, including the formation of large intracellular vacuoles highlighted by staining ER markers, increased actin fibre thickness, and disrupted Golgi and mitochondrial networks (Fig.9A-C). Mitochondrial respiration was also impaired, with reductions in both basal and maximal rates (Fig.9D-E). While mitochondrial activation based on MitoTracker intensity remained unchanged, TOM20 intensity was reduced, consistent with the findings in KALRN-deficient cells (Supplementary Fig.8C-E). TMCO1-deficient cells also showed decreased proliferation, as indicated by reduced Ki67 staining, and increased senescence by SAβG activity (Fig.9F-I). Comparison between TM cells from a young and an aged donor suggested an age-related difference in the extent of the senescence response (Supplementary Fig.8F-G). Strikingly, TMCO1 knockdown led to reduced expression of KALRN-12, further supporting the functional interaction between KALRN and TMCO1 (Fig.9 J-K; Supplementary Fig.8H).

**Figure 9:**
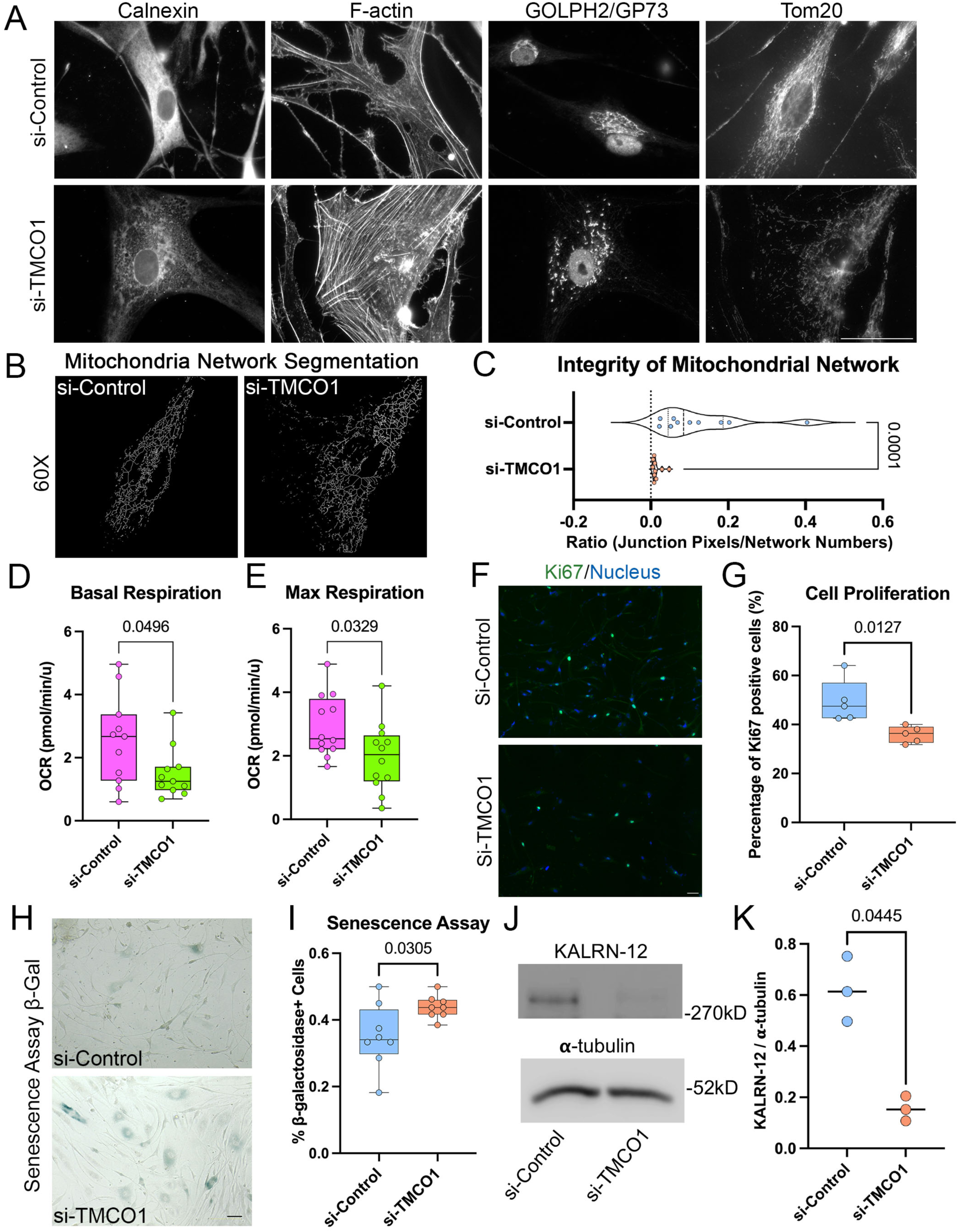
TMCO1 knockdown replicates phenotypic effects of KALRN depletion in human TM cells. **(A)** TMCO1 knockdown caused large vacuoles and abnormal structure of cytoskeleton, mitochondrial network, ER and Golgi in TM cells, shown by staining for the ER (calnexin), actin filaments (phalloidin), Golgi (GOLPH2/GP73) and mitochondria (TOM20). Scale bar, 50μm. **(B)** Segmentation of TOM20 positive mitochondria networks showed decreased network integrity in TMCO1 depleted cells. **(C)** Quantification of mitochondrial network structure revealed reduced junction pixel-to-network ratios (n=10 cells for each group, TM155; Mann Whitney test, fold change relative to control (FC)=0.12). **(D–E)** Mitochondrial respiration analysis by Seahorse assays showed reduced basal and maximal oxygen consumption rates in TMCO1-deficient cells (n=11-12 replicates per group; unpaired t-tests). **(F–G)** TMCO1 knockdown reduced cell proliferation, as shown by lower Ki67 (green) positive cells (n=5 images per group; unpaired t-test). Scale bar, 50μm. **(H-I)** β-galactosidase staining (blue) indicated increased senescence in TMCO1-depleted cells (n=8 images for each group; unpaired t-test). Scale bar, 50μm. **(J-K)** TMCO1 loss reduced expression of Kalirin-12 (MW: ∼470kDa; n=3 technical replicates; paired t*-*test).

**Figure 10:**
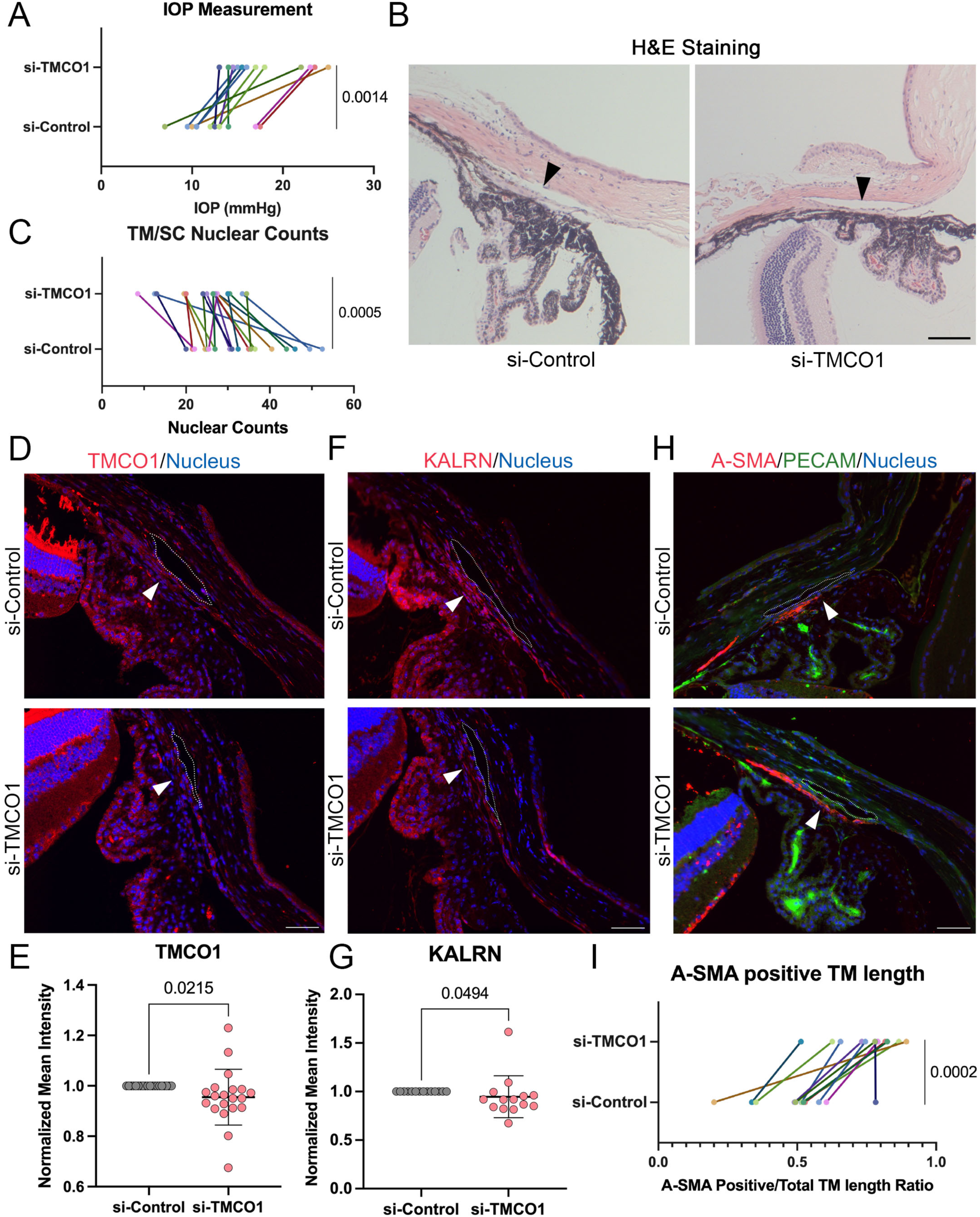
In vivo depletion of TMCO1 elevates intraocular pressure and reduces trabecular meshwork cellularity. **(A)** IOP was increased in eyes treated with TMCO1 siRNA compared to control eyes (n=12, paired t-test). **(B, C)** H&E staining showed reduced cell density in the TM, along with increased pigmentation in TMCO1-depleted tissue (n=20; paired t-test). Black arrows: Schlemm’s canal; Scale bar, 50μm. **(D, E)** Immunostaining confirmed knockdown of TMCO1 (red) in treated eyes (n=20; Wilcoxon matched-pairs signed-rank test). White arrow: TM tissue; dashed circle outlines SC. Scale bar, 50μm. **(F, G)** TMCO1 knockdown also reduced KALRN expression (red) in TM tissue (n=14; Wilcoxon matched-pairs signed-rank test). White arrow: TM tissue. Dashed circle outlines the SC. Scale bar, 50μm. **(H, I)** α-SMA staining (red) was more broadly distributed in the TM following TMCO1 knockdown (n=12, paired t-test). White arrow: TM tissue. Dashed circle outlines the SC. Scale bar, 50μm.

To assess the impact of TMCO1 knockdown on IOP, Accell siRNAs were injected into the anterior chambers, performing again a pairwise analysis by injecting control siRNAs into one eye and *TMCO1* siRNAs into the other one. Mice were then monitored by taking IOP readings from day 4 to 8. IOP values showed a significant increase following TMCO1-depletion compared to controls (Fig.10A). H&E staining revealed reduced nuclear density in the TM and SC inner wall in TMCO1-deficient tissue (Fig.10B-C). Immunostaining confirmed efficient TMCO1 knockdown in the treated eyes (Fig.10D-E). A concurrent reduction in KALRN expression was observed in TMCO1-depleted TM tissue (Fig.10F-G), consistent with their functional interaction observed *in vitro*. Similar to KALRN knockdown, TMCO1-depletion led to an expanded distribution of α-SMA-positive cells within the TM (Fig.10H-I).

Together, these findings demonstrate that TMCO1-depletion elevates IOP and induces structural changes in the TM that closely resemble those caused by KALRN suppression, supporting a shared functional pathway in TM homeostasis and reciprocal regulation of the two glaucoma risk genes.

## Discussion

We have identified the POAG-associated Rac1 GEF KALRN as a key regulator of TM cell physiology *in vitro* and *in vivo*. Our findings reveal that KALRN regulates TM cell homeostasis and IOP in concert with TMCO1. KALRN and TMCO1 regulate each other and depletion of either one leads to cell-wide changes such as ER and Golgi disruption, disturbed energy metabolism, and induction of cellular senescence. Depletion of either protein also led to increased IOP and glaucomatous changes in the TM *in vivo*. Thus, KALRN and TMCO1 are central regulators of the TM, highlighting functional interplay been genetically identified POAG-genes leading to disruption of molecular mechanisms central to cell homoeostasis and function and, thereby, increased IOP and disease.

Previous single-cell transcriptomic analyses of the anterior chamber identified KALRN as predominantly expressed in the ciliary muscle and TM cells^25^. In the present study, KALRN protein expression was similarly localized to these regions, but was notably higher in the TM compared to the ciliary muscle*. In vivo* depletion of KALRN in the TM led to a strong increase in IOP, suggesting that KALRN is of functional importance in the TM for regulating IOP.

KALRN-depletion induced cell-wide changes indicating that it regulates a signalling mechanism central to TM cell homeostasis. KALRN has two GEF domains one for Rac1/RhoG and one for RhoA. Complementation experiments indicated that the C-terminal RhoA GEF domain was not required to rescue the KALRN-depletion phenotype. A Rho GTPase Biosensor analysis revealed that Rac1 was strongly downregulated upon KALRN-depletion. In contrast, RhoA was induced. Hence, KALRN functions as a major Rac1 GEF, and not RhoA, activator in TM cells. Pharmacologic inhibition of Rac1 recapitulated key features of the KALRN-depletion phenotype, further corroborating the importance of Rac1 signalling downstream of KALRN. RhoA and Rac1 – or in some cells RhoA and Cdc42 – are often regulated in opposing manners^46–48^. RhoA activation upon KALRN-depletion and Rac1 downregulation is likely reflecting a more mesenchymal phenotype as α-SMA was also strongly induced and is likely to involve activation of downstream RhoA GEFs, such as GEF-H1^49–51^. KALRN-depletion also stimulated a profound reorganisation of microtubules from parallel arrays to radial arrays originating from a perinuclear centre, commonly observed in mesenchymal cells, which, in turn, may impact on mitochondria as well as ER and Golgi structure, distribution, and function^52^. Rac1 is a major regulator of the microtubule cytoskeleton; hence, the reduced Rac1 activity is likely to impact microtubule organisation and thereby promote the observed downstream RhoA activation and F-actin reorganisation^53^.

KALRN-depletion induced large vacuoles containing luminal soluble ER markers and secretory cargo leading to reduced expression of ER-associated proteins important for ER function, such as BiP and TMCO1. Rac1 is an important regulator of microtubule/ER dynamics, calcium signalling, and secretory trafficking via COPII vesicles^54, 55^; hence, KALRN as an ER-associated Rac1 GEF impacts on both microtubule organisation and ER function. ER dysfunction and disruption of calcium homeostasis is likely to lead to other KALRN-depletion induced phenotypes such as inhibition of autophagy as the ER provides lipids to forming autophagosomes, and disruption of the Golgi apparatus as Rac1 stimulates COPII vesicle trafficking and inhibition of COPII vesicle formation^41, 55–57^.

KALRN-depletion resulted in a fragmented mitochondrial network accompanied by increased DRP1 expression, indicating enhanced mitochondrial fission. The mitochondrial import receptor TOM20 was strongly downregulated, indicating a reduction in mitochondrial mass; hence, oxidative phosphorylation was strongly reduced in KALRN depleted cells. Mitochondria rely on ER-Mitochondria contact sites for lipid and calcium transfer; hence, the dysfunctional ER and disrupted calcium homeostasis caused by KALRN-depletion likely cause the loss of mitochondrial mass and, hence, reduced oxidative phosphorylation^58^. As ER-Mitochondria contact sites are lost by aging cells, KALRN malfunction may accelerate the normal aging process^59^. This is supported by the observation that KALRN-depletion led to a strong increase in cellular senescence.

In POAG patients, TM cellularity is markedly reduced, accompanied by an increase in senescent TM cells compared to age-matched non-glaucomatous controls^1, 60^. In the present study, we demonstrate that depletion of KALRN induces cellular senescence and reduces TM cell density *in vivo*. Interestingly, KALRN was also reported to be increased in TM progenitor cells and has been identified through bioinformatic analyses as a potential hub gene involved in the regulation of TM progenitor cell development^61^. These findings suggest that KALRN plays a critical role in maintaining TM cell homeostasis by preventing entry of mature TM cells into cellular senescence and supporting TM progenitor cell function.

Abnormal TM phenotypes are a hallmark of POAG, characterized by a shift toward a fibrotic state. Glaucomatous TM cells display elevated α-SMA^62–64^, thick F-actin stress fibres, and increased cross-linked actin networks (CLAN)^64–66^. Inactivation of some POAG genes has recently been shown to alter the actin cytoskeleton^67^. The TM cells with increased CLANs were shown to be stiffer which may contribute to the disease pathology. In this study, increased α-SMA positive TM cells, thicker F-actin stress fibres, and increased CLAN-like structures were observed in KALRN depleted TM cells.

A key cytoskeletal marker of the fibrotic shift in glaucomatous TM cells is α-SMA, an actin isoform incorporated into stress fibres that enhances cellular contractility and extracellular matrix remodelling. In vitro, α-SMA expression is notably induced by TGF-β1/2 - growth factors elevated in the aqueous humour of POAG patients and widely used to model the disease^68–71^. Single-cell transcriptomic profiling has revealed that α-SMA is predominantly expressed in a specific TM subpopulation, termed TM-subtype 3, which localizes to the inner uveal meshwork and may be selectively expanded under glaucomatous conditions^72^. In our study, KALRN-depletion led to a marked increase in α-SMA-positive TM cells, along with thicker F-actin stress fibres and increased CLAN formation - hallmarks of glaucomatous remodelling. However, despite these features, KALRN-depleted cells exhibited reduced contractility and motility, likely due to the concurrent drop in energy metabolism caused by the dysfunction of the ER. The enrichment of α-SMA-positive cells may reflect a shift toward the TM-subtype 3 identity, which has also been predicted to be uniquely affected by pronounced mitochondrial pathway changes^72^. These findings raise the possibility that KALRN modulates both cytoskeletal dynamics and energy metabolism in distinct TM cell subtypes, a relationship that merits further investigation.

In the anterior chamber, melanin granules shed from the iris pigment epithelium and ciliary body disperse into the aqueous humour as part of normal physiological turnover^73^. TM cells internalize this pigment through phagocytosis to maintain outflow function and anterior segment homeostasis^73^. In pigment dispersion syndrome (PDS), excessive pigment accumulation in the TM leads to cellular loss and elevated IOP^73^. In our study, KALRN-depletion resulted in increased pigmentation in TM tissue, suggesting that KALRN-depletion disrupts pigment turnover in trabecular meshwork cells, possibly due to the disrupted endo-lysosomal system, highlighting a potential regulatory mechanism that warrants further investigation.

Depletion of either KALRN or TMCO1 resulted in strikingly similar cellular phenotypes, characterized by prominent cytoplasmic vacuolization, enhanced F-actin bundling, and disruption of both Golgi and mitochondrial architecture. Silencing of one gene led to reduced expression of the other, indicating reciprocal regulation. As both proteins are associated with the ER, reduced expression may be the result of ER disruption caused by depletion of either one of the two proteins. The convergence of phenotypic outcomes and co-expression patterns strongly supports a synergistic role for KALRN and TMCO1 in maintaining cellular homeostasis; hence, this synergy may amplify small effects caused by genetic variants in the two genes and, thereby, contribute to the complex genetics of POAG.

Our study highlights KALRN as a critical regulator of trabecular meshwork cell homeostasis, as its depletion disrupts ER and secretory organelle structure, stimulates cytoskeletal remodelling, and disrupts energy metabolism, leading to elevated IOP. Our results also identify that depletion of TMCO1, a gene at one of the most strongly POAG-linked loci, produces similar phenotypes and its expression is reciprocally linked to KALRN. Thus, our study demonstrates that two genes linked to increased IOP and POAG in patients due to their proximity to risk loci for glaucoma indeed regulate IOP in an interdependent manner. A large number of genes has been linked to high IOP and POAG by studies investigating the complex genetics of glaucoma. Exploiting that knowledge to identify the functional links between the KALRN/TMCO1 module and other glaucoma genes, and to discover additional regulatory modules will be essential to advance targeted therapeutic strategies aimed at preserving trabecular meshwork function and preventing glaucoma progression

## Methods

### TM Cell Culture

Four human trabecular meshwork (TM) cell strains (Supplementary Table 1) derived from different donors were isolated and authenticated according to consensus recommendations as previously described^74, 75^. The TM cells were cultured in low glucose Dulbecco′s Modified Eagle′s Medium (DMEM) (Sigma-Aldrich, USA) supplemented with 10% Fetal Bovine Serum (FBS, Sigma-Aldrich, USA), 1% Penicilin-Streptomysin (Pen/Strep, Sigma-Aldrich, USA) at 37°C in a 5% CO₂ incubator. The medium was changed every other day until 80% confluence. The cells from passages 4-7 were used in the experiments. For immunocytochemistry, TM cells were plated on coverslips pre-coated with 10% Matrigel (Corning®, UK).

### SiRNA Transfection

Cells were transfected with siRNAs (Merck, USA) using Lipofectamine™ RNAiMAX (Invitrogen, USA) following the manufacturer’s protocol. Briefly, siRNAs (20µM stock) and RNAiMAX reagent were separately diluted in Opti-MEM™ I Reduced Serum Medium (Giboco, UK) at a 1:25 ratio and then combined and incubated for 15 minutes at room temperature to allow complex formation. The siRNA-RNAiMAX complexes were further diluted in TM culture medium containing 10% FBS, 1% Pen/Strep, and 100μM HEPES buffer (Sigma-Aldrich, USA) at a 1:5 ratio, resulting in a final siRNA concentration of 80nM. Cells were incubated with the transfection mixture for 24 hours. Subsequently, the medium was replaced with fresh TM culture medium containing 10% FBS, and cells were cultured for an additional 48 hours before further experiments. The following human siRNAs were used: *si-Control*: 5′-UGGUUUACAUGUCGACUAA-3′, 5′-UGGUUUACAUGUUGUGUGA-3′; *si-KALRN*: 5′-GCUACAUUGUCAACCGGGU-3′, 5′-CAUAUCAUCUUUGGCAACA-3′, 5′-CUCUAGAUGCCACACUCAA-3′; and *si-TMCO1*: 5′-GGACAGACAAGUACAAGAG-3′, 5′-GAAGAGAAACUGAAGAAUA-3′. If not otherwise indicated, experiments were performed with pools of siRNAs. Independent of using pools or individual siRNAs, the total concentration was kept constant. All siRNAs were synthesized with dTdT 3’-overhangs.

### Plasmids

The expression plasmids for rat KALRN isoforms pEAK10-His-Myc-Kal7 and pEAK10-His-Myc-Kal12 were gifts from Betty Eipper (Addgene plasmids # 25454; http://n2t.net/addgene: 25454; RRID:Addgene_25454 and # 25442; http://n2t.net/addgene:25442; RRID: Addgene_25442) ^76, 77^. The RhoA biosensor expression plasmid pCAGGS-Raichu-RhoA-CR (Addgene plasmid # 40258; http://n2t.net/addgene:40258; RRID:Addgene 40258) and pcDNA3-Clover (Addgene plasmid # 40259; http://n2t.net/addgene:40259; RRID: Addgene_40259) were gifts from Michael Lin^44^. pCAGGS-Raichu-Rac1-CR was constructed by replacing the sequence between Clover and mRuby2 in pCAGGS-Raichu-RhoA-CR with the corresponding sequence of pCAGGS-Raichu-Rac1 (kindly provided by Michiyuki Matsuda; Department of Pathology and Biology of Diseases, Kyoto University, Japan). The sequence of the resulting plasmid was confirmed by Nanopore sequencing.

### DNA Transfection of TM cells

TM cells were transfected using Lipofectamine LTX (Invitrogen, USA) in a 48-well format, following the manufacturer’s protocol. One day prior to transfection, cells were seeded at a density of 1 × 10⁴ cells per well in 200μl of TM cell culture medium to achieve 50–80% confluency. For each well, 0.25μg of plasmid DNA was diluted in 50μl of Opti-MEM I (Gibco™, USA) and combined with 0.25μl of Lipofectamine PLUS Reagent. After a 15-minute incubation, 1μl of Lipofectamine LTX was added, mixed thoroughly, and incubated for an additional 25 minutes to facilitate complex formation. The transfection mixture (50μl) was then added dropwise to each well following the addition of 250μl of fresh TM cell culture medium. Cells were incubated at 37°C for 4 hours, after which transgene expression was assessed. Reagent volumes were adjusted accordingly for different plate formats as per the manufacturer’s recommendations.

### Drug treatments

The Rac1 inhibitor NSC 23766, Bafilomycin A1, and Thapsigargin were purchased from Bio-Techne/Tocris. NSC 23766 was dissolved at 100mM in DMSO and then diluted to 100μM in tissue culture medium. Bafilomycin A1 was dissolved at 100µM in DMSO and diluted to 200nM with tissue culture medium. Thapsigargin was dissolved at 5mM in DMSO, and the cells were then treated with 5µM diluted in Live Cell Imaging Solution.

### Calcium Imaging Assay

TM cells were washed twice with fresh TM culture medium containing 10% FBS and incubated with 2 µM Fluo-4 AM (Invitrogen, CA), a fluorescent calcium indicator, for 1 hour at 37°C in the dark. Following incubation, the cells were washed twice with Live Cell Imaging Solution (Gibco™, USA) and maintained in Live Cell Imaging Solution during live-cell imaging. Fluo-4 fluorescence (excitation at 488 nm) was captured using a Nikon Ti-E Live Cell Microscope (Nikon, Japan) and a CFI Apochromat Nano-Crystal 60x oil objective (N.A., 1.2). Subsequently, the cells were treated with 5 µM Thapsigargin for 10 minutes to induce net calcium release into the cytosol. Post-treatment Fluo-4 fluorescence was imaged under the same conditions. Fluorescence intensity was quantified using ImageJ-Fiji software ^78^, with at least 20 cells analysed per group to assess differences in intracellular calcium levels between control and *KALRN* siRNA-treated cells.

### Contraction Assay

siRNA-transfected TM cells were harvested and prepared for collagen gel contraction assays. Cells were trypsinized for 2–5 minutes at 37°C until detachment could be observed by microscopy. The trypsin was inactivated with fresh TM culture medium, and the cell suspension was collected and adjusted to a final cell number of 1 × 10⁵ cells per gel, with an additional 10% added to account for potential cell loss. An extra 5 mL of fresh medium was added to the suspension, which was followed by centrifugation at 1000 rpm for 5 minutes. The resulting cell pellet was resuspended in 100μL FBS for subsequent use. Collagen gels were prepared by combining 1000μl rat tail collagen type I (2.05 mg/mL) with 160μl of concentrated medium (7.4% L-glutamine, 2mM, Sigma-Aldrich, USA and 18.9% Sodium Bicarbonate, 7.5% solution, Gibco™, USA in 10x Dulbecco′s Modified Eagle′s Medium - low glucose, Sigma-Aldrich, USA)). The pH was adjusted with 1M NaOH until the solution turned stable salmon pink, indicating the correct pH for polymerization. The prepared cell suspension was then added to the collagen mixture, and 150μL of the final gel solution was cast into the wells of 35 mm MatTek tissue culture dishes (MatTek Corporation, Ashland, USA) (3 replicates per experimental group), ensuring even distribution without air bubbles. Gels were polymerized at 37°C for 10 minutes (leftover gel in the tube was used as a polymerization control). To generate relaxed gels, the edges were detached using a pipette tip after polymerisation, and any unpolymerized solution was carefully removed. Each gel was overlaid with 2ml growth medium and incubated at 37°C. The medium was refreshed every 2 days. Gels were monitored daily, and images were captured for subsequent analysis. For quantifying gel contraction, the areas of the gels and corresponding MatTek wells were measured using the Image J-Fiji ^78^. The contraction was calculated using the following formula: 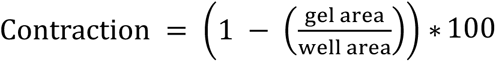.

### Live Cell Imaging, Cell Movement Quantification, and FLIM FRET

For cell migration analysis, 1×10^4^ cells were plated per well in a 24-well plate (three replicates). Following transfection with siRNAs, cells were incubated in fresh culture medium containing 10% FBS for 48 hours. Cells were then washed twice with Live Cell Imaging Solution (Gibco™, USA) supplemented with 10% FBS and 1% Pen/Strep and maintained in 1 mL of the same solution. Time-lapse imaging was performed using a Nikon Eclipse Ti-E microscope equipped with a Nikon S Plan Fluor 20×objective (N.A. 0.45) at 37°C. Images were acquired every 2 minutes, capturing 63 frames per field of view across six distinct areas per group. Cell movement was quantified based on two parameters: the curved trajectory distance and the similarity of cell shape between the initial and final timepoints. To measure trajectory distance, cells were segmented at frames 1, 10, 20, 30, 40, 50, and 60 using Image J/Fiji. The centre of segmented cells was measured by using a previously described Python 3.8.0 (https://www.python.org/) algorithm ^79^. Cell centre trajectories and corresponding curved distances from frame 1 to 60 were visualized and analysed using a previously described Python (v 3.8.0) algorithm ^79^. To quantify cell shape similarity between the initial and final time points, cells in frames 1 and 63 were segmented. The cell centres in both frames were identified and aligned. Shape similarity was assessed using the Hausdorff Distance, computed with a previously described Python algorithm ^80^ to compare segmented cell shapes between frames 1 and 63. A higher Hausdorff Distance indicates lower shape similarity. To analyse activity of RhoA and Rac1 by fluorescence lifetime-based Foerster resonance energy transfer microscopy (FLIM FRET) was performed using a Leica Stellaris 8 confocal microscope and 63× (N.A., 1.40) oil objective and the FLIM module of the LAS X software (Leica Microsystems, Wetzlar, Germany). Cells were transfected with either pCAGGS-Raichu-RhoA-CR, pCAGGS-Raichu-Rac1-CR, or pcDNA3-Clover. Cells were analysed in Live Cell Imaging Solution at 37°C. Images of the pcDNA3-Clover transfection were used to determine the lifetime of the donor. Images were analysed using the Phaser module of the Leica LAS X single molecule detection (Falcon) software.

### Mitochondria Activity Assay

Cells were stained with MitoTracker Orange CMTMRos (M7510; ThermoFisher Scientific) following the manufacturer’s guidelines with some modifications. A 1 mM stock solution was prepared by dissolving 50 µg MitoTracker Orange CMTMRos in 117 µL DMSO. To improve solubility, the stock solution was diluted in PBS (1:50) to prepare an intermediate working solution (stock-2). For staining, stock-2 was added to Seahorse XF DMEM Medium, PH7.4 (1:100; Agilent, USA) to achieve a final concentration of 200nM. Cells were incubated with 200 µL of the staining solution per well in a 48-well plate at 37°C for 1 hour. Following incubation, cells were fixed with 3% paraformaldehyde (PFA) for 20 minutes at room temperature, permeabilized for 5 minutes, and blocked in blocking buffer for 1 hour. Cells were stained with TOM20 primary antibody (1:100, Santa Cruz Biotechnology, USA) overnight at 4°C, followed by incubation with Donkey anti-Mouse IgG Alexa Fluor™ 488 secondary antibody for 2 hours at room temperature. Samples were mounted and imaged using a Nikon Eclipse Ti-E microscope equipped with a Nikon S Plan Fluor 20× objective (N.A. 0.45). Staining intensities of TOM20 and MitoTracker were quantified using ImageJ/Fiji, and the MitoTracker/Tomm20 ratio was calculated as a measure of mitochondrial activity.

### Seahorse Assays

Cells were seeded into a Seahorse XFe96/XF Pro Cell Culture Microplate (Agilent, USA) at a density of 0.7*10^4^ cells/well in 200ul TM culture medium. After cultured in the fresh TM culture medium for 24 hours, the cells were transfected with siRNAs for 24 hours followed by incubating in the fresh culture medium for 48 hours. One hour prior to the Seahorse assay, the culture medium was replaced with Seahorse XF Assay Medium, composed of Seahorse XF DMEM Medium (pH 7.4, Agilent, USA) supplemented with 10 mM glucose, 1 mM pyruvate, and 2 mM L-glutamine (Agilent, USA). All experiments measuring energy metabolism were run with a Seahorse XF Pro Analyzer using the software to run and analyse the experiments provided by the manufacturer (Agilent, USA). The mitochondrial stress test, the glycolytic rate test, and the ATP rate tests were run according to the manufacturer’s instructions using the reagents provided with the respective kits for these assays or components purchased from Merck Lifescience UK (rotenone, R8875; antimycin A, A8674; FCCP, C2920; oligomycin, 495455; and 2-deoxyglucose, D6134). Assay results were normalized by quantifying cell numbers using the CyQUANT assay. Briefly, cells in the Seahorse Cell Culture Microplates were frozen overnight at −80°C. The CyQUANT assay was performed using the CyQUANT™ Cell Proliferation Assay Kit (Invitrogen, USA), following the manufacturer’s guidelines. Fluorescence was measured at an excitation wavelength of 480 nm and an emission wavelength of 520 nm. The CyQUANT data were uploaded into the Seahorse Wave software for normalisation of the Seahorse experiments.

### Senescence-associated β-galactosidase Activity Assay

Cells were cultured under appropriate conditions and fixed with 3% paraformaldehyde (PFA) in PBS for 5 minutes at room temperature. For senescence-associated β-galactosidase (SAβG) staining, cells were incubated overnight at 37°C (without CO₂) in SAβG staining solution containing 0.1% X-gal (5-bromo-4-chloro-3-indolyl β-D-galactopyranoside; Roche/MERCK, Germany), 5 mM potassium ferrocyanide, 5 mM potassium ferricyanide, 150 mM sodium chloride, 2 mM magnesium chloride, and 40 mM citric acid/sodium phosphate buffer (pH 6.0), with pH 4.0 and 7.5 serving as positive and negative controls, respectively. Following incubation, the staining solution was removed, and cells were washed three times with distilled water. The cells were then stained with Hoechst (1:2000 diluted in distilled water) to label nuclei. Samples were maintained in distilled water and imaged using an Invitrogen EVOS XL Core microscope or a Nikon Eclipse Ti-E microscope. SAβG activity was quantified by determining the proportion of SAβG-positive (blue, bright-field) cells at pH 6.0 relative to the total nuclear count (Hoechst fluorescence) per field.

### Autophagy Flux Assessment

To measure autophagic flux, cells were treated with Bafilomycin A1 (200nM, diluted in culture medium; TOCRIS, UK) for 6 hours. Following treatment, cells were washed twice with sterile PBS, and total protein was extracted for LC3-I/II quantification by immunoblotting. Autophagic flux was determined by calculating the difference in LC3-II expression between Bafilomycin A1 treated (Baf⁺) and untreated (Baf⁻) conditions and comparing these values between control and *KALRN* knockdown TM cells.

### Immunostaining and Fluorescence Microscopy

Cell fixation and processing for immunofluorescence were performed as previously described ^81, 82^. Briefly, cells were fixed using either methanol for 5 minutes at −20°C, followed by rehydration in PBS at room temperature, or 3% PFA in PBS for 20 minutes at room temperature. PFA-fixed cells were permeabilized with 0.3% Triton X-100 (Sigma-Aldrich, USA) in PBS containing 0.5% BSA (Capricorn, UK)and 20 mM glycine (Sigma-Aldrich, USA) for 3 minutes, followed by two washes with blocking buffer (PBS with 0.5% BSA and 20 mM glycine ^81^). The samples were incubated in blocking buffer for 1 hour, followed by overnight incubation with primary antibodies Table. The samples were then washed twice with blocking buffer and incubated with the secondary antibodies for 2 hours. After two washes, samples were embedded in ProLong Gold antifade reagent (Life Technologies, USA). Imaging was conducted using a Nikon Eclipse Ti-E microscope with a Nikon S Plan Fluor 20× objective (N.A., 0.45) or 60× oil objective (N.A., 1.4) lens (Nikon Europe, Amstelveen, The Netherlands) or a Leica TCS SP8 microscope with a 63× (N.A., 1.40) oil objective (Leica Microsystems, Wetzlar, Germany). Images were processed using ImageJ/Fiji.

### Immunoblotting

Immunoblotting was performed as described before ^82^. Briefly, whole-cell lysates were collected by washing cells with PBS, followed by the addition of SDS-PAGE sample buffer and heating at 70°C for 10 min. The samples were homogenized using a 1 ml syringe with a 25G needle, and proteins were separated by SDS-PAGE before being transferred onto PVDF membranes ^83^. Membranes were blocked with 5% defatted milk powder dissolved in TBS containing 0.1% Tween-20 and incubated overnight with primary antibodies in the same blocking solution. For anti-phospho-protein antibodies, blocking and incubation were performed using 5% BSA in TBS with 0.1% Tween-20. After extensive washing, membranes were incubated with HRP-conjugated or fluorescent secondary antibodies. Protein detection was carried out using either the Bio-Rad ChemiDoc ECL detection system (Bio-Rad Laboratories, Hercules, CA, USA) or the Li-Cor ODYSSEY infrared imaging system (Li-Cor, Lincoln, NE, USA). The antibodies and related dilution used for Immunoblotting are shown in the Table.

### siRNA Injection of Mouse Eyes and IOP Measurements

Accell smart pool siRNAs were purchased from Horizon Discovery Biosciences Limited (Cambridge, UK; KALRN, E-047265; TMCO1, E-044628; Non-targeting, D-001910). The efficacy of Dharmacon™ Accell™ siRNAs was first validated *in vitro* using the mouse kidney cell line IMCD (kindly provided by Paul Gissen, UCL, UK). Cells were transfected with Accell siRNAs following the protocol provided by the manufacturer for transfection without transfection reagents. Adult C57BL/6J mice (10–12 weeks old) were anesthetized by intraperitoneal injection of Ketamine/Dormitor (30ug/5ug per g of body weight) and pupils were dilated using topical mydriatic solution (Pharma Stulln, #ATH0713). 4 µl of Accell siRNAs (100µM) were then injected into anterior chambers essentially as described previously ^84^. Each mouse was injected with non-targeting control siRNA in one eye and an siRNA targeting either *KALRN* or *TMCO1* in the other eye, alternating left/right. Eyes were injected through the peripheral cornea under a surgical microscope using a 10 mm 34-gauge hypodermic needle mounted on a 5 μl syringe (Hamilton AG, Bonaduz, Switzerland). Prior to injection, a paracentesis was performed during which aqueous humour outflow could be observed. At the end of the procedure, Antisedan (atipamezole hydrochloride 0.10 mg/ml) was injected intraperitoneally, and all animals received chloramphenicol 1% eye ointment to the cornea. IOP was then measured from day 4 after injection using rebound tonometry (TonoLab, Icare) as previously descripted ^85^. Five consecutive measurements were taken per eye, each consisting of an average of six readings, and averaged for each reading. Eyes were then collected for further analyses by enucleation and overnight fixation in 10% Buffered Formalin (NBF, Sigma-Aldrich, USA). All experiments conducted were in accordance with the UK Home Office Animals Scientifics Procedures Act 1986 (PPL PP6039705).

### Immunohistochemistry

Formalin-fixed eyes were embedded in paraffin and 4μm sections were cut. Slides were deparaffinized by sequential incubation in Histoclear II (3 × 8 min; National Diagnostics, USA) followed by rehydration through graded ethanol solutions (100% ethanol: 2 × 6 min; 95%, 90%, 80%, and 70% ethanol: 5 min each) and MilliQ water (5 min). Antigen retrieval was performed using Citrate Antigen Retrieval Buffer (pH=6, containing 8.25mM sodium citrate and 1.75mM citric acid). The buffer was preheated in a water bath at 95°C before immersion of the slides. Slides were incubated in the retrieval buffer for 35 min and then cooled at room temperature for 5 min before rinsing with PBS. The slides were then stored submerged in PBS at 4°C until staining. For immunostaining, sections were incubated with primary antibodies (Table) diluted in PBS-0.3% Triton-X (Sigma-Aldrich, USA), to enhance antigen penetration. Approximately 150μl of the diluted antibody solution was applied per slide. Following primary antibody incubation, slides were washed three times for 10 minutes each in prewarmed PBS-0.1% Tween20. Secondary antibodies (Table) were then applied at a dilution of 1:500 in 0.3% Triton-X, along with Hoechst (1:500) for nuclear counterstaining. After incubation, slides were washed three times in 0.1% PBS-Tween for 10 minutes each, followed by a final wash in PBS for 10 minutes. The slides were then mounted using ProLong™ Gold Antifade Mountant (Invitrogen, USA). The staining was imaged using a Nikon S Plan Fluor 20× objective (N.A., 0.45) or 60× oil objective (N.A., 1.4) lens (Nikon Europe, Amstelveen, The Netherlands) or a Leica TCS SP8 microscope with a 63× (N.A., 1.40) oil objectives (Leica Microsystems, Wetzlar, Germany). The staining intensity was analysed using ImageJ/Fiji.

### Image Analysis

#### ZO1 distribution quantification

To assess ZO-1 distribution in cells, immunofluorescence staining was performed using a ZO-1 antibody, followed by imaging with a Nikon Eclipse Ti-E microscope equipped with a Nikon S Plan Fluor 20× objective lens (N.A. 0.45). For each experimental group, 5–6 images were acquired. Cells were categorized into three distinct types based on ZO-1 distribution: Type-1, cells fully enclosed by continuous ZO-1 staining; Type-2, cells partially enclosed by continuous ZO-1 staining extending over more than half of the cell’s long axis; and Type-3, cells minimally enclosed or lacking continuous ZO-1 staining (covering less than half of the long axis). The number of cells in each category was quantified using the Cell Counter plugin in ImageJ/Fiji ^86^, and the percentage of each cell type was calculated.

#### Cell Shape

To validate the cell shape changes, cells (n>400) stained with ZO-1 antibody and Hoechst were segmented using the Cellpose algorithm ^87^. To enhance segmentation accuracy, a custom Cellpose model was trained using manually annotated images of 50 cells per treatment group. The segmentation were uploaded to VAMPIRE software ^88^ for cell shape analyses. The contour of each individual cell was represented by 50 equidistant points along its perimeter. To enable comparative and fair analysis, cell shapes from a given dataset were pooled, normalized for size, and aligned. Principal component analysis (PCA) was then applied to the contour coordinates of the aligned cells to derive eigenshape vectors (i.e., principal components, PCs). Cell shapes were reconstructed using a reduced set of eigenshape vectors, with the number of vectors selected to account for 95% of the total shape variance across all analysed cells. To identify representative cell shape modes, K-means clustering was applied to the morphological data described by the reduced eigenshape vectors. The percentages of each cell shape cluster were quantified and represented in a heatmap.

#### Integrity of Mitochondria and Golgi Network

To assess mitochondrial and Golgi network integrity, cells were stained with the mitochondrial marker TOM20 or a Golgi marker and imaged at 20× magnification. Network segmentation and quantification were performed using ImageJ/Fiji, following previously established protocols ^89^. Mitochondrial and Golgi networks from over 20 cells were segmented using the ROI Manager tool in ImageJ. To enhance network features, the Unsharp Mask, Enhance Local Contrast (CLAHE), and Median Filter functions in ImageJ/Fiji were applied. The networks were then binarized and skeletonized using the Make Binary and Skeletonize tools. Structural parameters, including network junction pixels, branch numbers, and branch lengths, were quantified using the Analyse Skeleton tool ^89^. To quantify network integrity, the ratio of junction pixels to branch number and branch length to cell area were calculated.

#### Cytoskeleton Structure Quantification

To analyse the structure of phalloidin-stained F-actin and anti-α-tubulin-stained microtubules, we utilized FSegment ^90^ and SOAX software ^91^, respectively. For F-actin analysis, cells were stained with phalloidin and imaged using a Nikon Eclipse Ti-E microscope equipped with a Nikon S Plan Fluor 40× objective lens (N.A. 0.6). A total of 20 cells were imaged per treatment group. Single-cell segmentation was performed using the ImageJ/Fiji selection tool and ROI manager. The segmented cytoskeletal structures were converted to grayscale and uploaded into FSegment for quantification. The density of thin (1–2 pixels in width) and thick actin filaments (3–15 pixels in width) was assessed, where 1 pixel corresponds to 0.16μm. For microtubule analysis, cells were stained with anti-α-tubulin and imaged using the same Nikon Eclipse Ti-E microscope and 40× objective lens. A total of 20 cells per treatment group were analysed. Single-cell segmentation was performed using ImageJ/Fiji and uploaded to SOAX for microtubule orientation (θ) and curvature analysis. To quantify microtubule disorder, the entropy of θ was calculated using the following formula: H(Θ) = −∑p(θ)log2p(θ), where p(θ) is the probability distribution of microtubule orientation, calculated as: 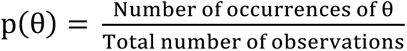. If H(Θ)=0, there is no disorder. If H(Θ) is high, there is more disorder.

#### Quantification of α-SMA-Positive Trabecular Meshwork Length

To analyse the α-SMA-positive trabecular meshwork (TM) length, paraffin-embedded mouse eye sections were stained with an α-SMA antibody and Hoechst. Imaging was performed using a Nikon Eclipse Ti-E microscope equipped with a 20× objective lens. Image analysis was conducted using ImageJ/Fiji. The total TM distance was defined as the distance from the distal end of Schlemm’s canal to the starting point of the TM, which was identified as the region beginning two cell layers away from the corneal endothelium’s single-cell layer. The α-SMA-positive TM length was defined as the distance from the distal end of Schlemm’s canal to the terminal boundary of α-SMA-positive trabecular meshwork. The ratio of α-SMA-positive TM length to the total TM length was then calculated to assess the relative distribution of α-SMA expression within the TM tissue.

#### Co-localization of a-SMA staining and Pigmentation

To assess the colocalization of α-SMA staining and pigmentation, mouse eye sections from siRNA-injected mice (5–8 days post-injection) were stained with α-SMA and Hoechst, then imaged using a Nikon Eclipse Ti-E microscope equipped with a 20× objective lens and fluorescence and bright-field channels. The TM region, defined as the area extending from the distal end of Schlemm’s canal to the initial boundary of the TM, was cropped for further analysis. To quantify the colocalization between α-SMA staining and pigmentation, the cropped TM images from the α-SMA fluorescence and bright-field channels were processed using the threshold function in Image J/Fiji to define positive α-SMA staining and pigmentation areas. The thresholded images were analysed using the IACoP plugin in ImageJ to calculate the Pearson’s correlation coefficient (r) and Manders’ overlap coefficients (M1: fraction of α-SMA overlapping pigmentation; M2: fraction of pigmentation overlapping α-SMA).

#### TM&SC Tissue Nucleus Counting

To evaluate the effect of gene knockdown on TM cell numbers in mouse TM tissue, the nuclei within the TM and Schlemm’s canal (SC) regions were quantified in H&E-stained sections using ImageJ/Fiji. Briefly, paraffin-embedded eye sections from control and gene knockdown mice were stained with Hematoxylin and Eosin (H&E) using a Leica ST 5010 Autostainer XL (Leica, Germany). Nuclei were counted within a defined region spanning from the distal end of Schlemm’s canal to the initial boundary of the TM, with the counting area restricted to the inner layer of Schlemm’s canal and the primary TM region. Nuclei quantification was performed using the Cell Counter plugin in ImageJ/Fiji.

#### Statistical Analysis and Reproducibility

Four different TM cell strains derived from 4 different donors were analysed. Quantifications of such data indicate the values for each strain and were analysed using paired tests. For quantifications of specific responses (e.g., Rho GTPase activity), data points of values derived from individual cells are shown. Such datapoints are derived from at least two independent cultures. Statistical significance was tested using Kruskal-Wallis and Wilcoxon tests, or ANOVA and t-tests. For pairwise multiple comparisons, Tukey HSD and Steel-Dwass tests were used. When not already part of the test, two-tailed tests were performed. All quantifications show all data points along with median and interquartile ranges or means and standard deviations. Graphs and statistical calculations were generated with JMP-Pro or GraphPad Prism.

The information of reagents and kits used in this study are shown in the Supplementary Tables.

## Data availability

The datasets generated and/or analysed during the current study are available from the corresponding author upon reasonable request.

## Acknowledgements

This work was supported by Moorfields Eye Charity (GR001476), the BBSRC (BB/X000575/1 and BB/Z516375/1), the MRC (MR/W028735/1), the NIHR Biomedical Research Centre at Moorfields Eye Hospital, and the UCL Institute of Ophthalmology. APK is supported by a UK Research and Innovation Future Leaders Fellowship (MR/Y033930/1), an Alcon Research Institute Young Investigator Award and a Lister Institute for Preventive Medicine Award.

## Author contributions

The study was conceived and planned by XF, APK, MSB, and KM with the support of DRO, WDS, JWBB, and MB. Most of the experiments were performed and analysed by XF with the support of KM, MSB, and JWBB. ERT and DRO supported the IOP measurements, and MB the contraction assays. WDS isolated the TM cell strains. XF wrote the manuscript with the support of all authors.

