## Supplementary Figures and Tables for "Interdependent regulation of trabecular meshwork cell physiology and intraocular pressure by KALRN and TMCO1"

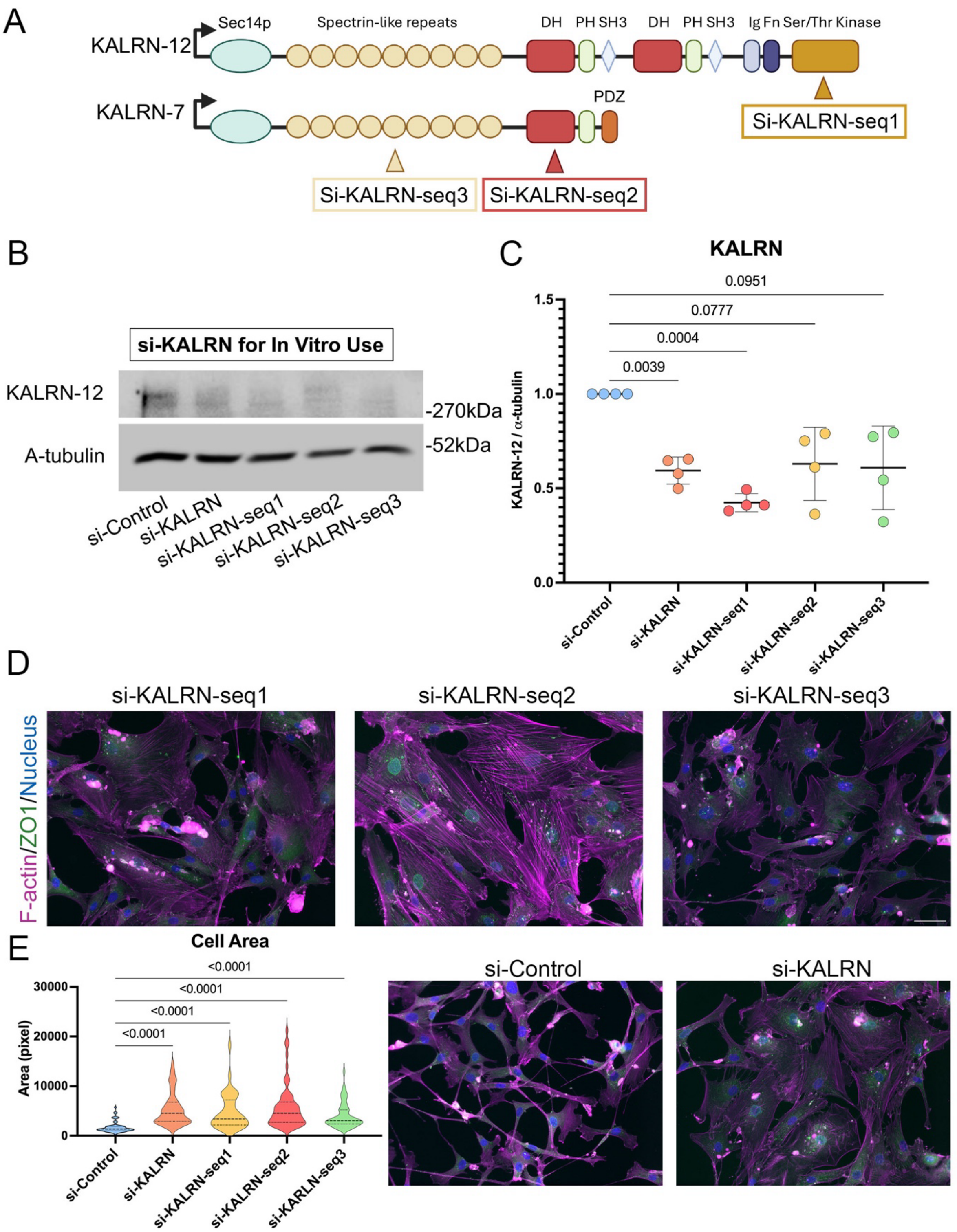

**Supplementary Figure 1: Targeted KALRN domains and efficiency of KALRN siRNAs.**

**(A)** Schematic of human KALRN isoforms, with domains and the specific target sites of the three distinct siRNAs (si-KALRN-seq1, -seq2, and -seq3) used in this study. The isoform and domain schematic was previously published[44]. **(B-C)** Western blotting showing efficient depletion of KALRN-12 (~340kDa) by individual siRNAs or their pool (n=4 replicates; RM one-way ANOVA). **(D)** TM cells transfected with individual or pooled KALRN siRNAs show similar cell morphology and comparable redistributions of ZO-1 (green) and F-actin (purple) by immunostaining. Scale bar, 50 µm. **(E)** Quantification of cell area shows an increase in TM cells transfected with individual or pooled KALRN siRNAs (si-control: n=72; si-KALRN pool: n=53; si-KALRN-seq1: n=41; si-KALRN-seq2: n=58; si-KALRN-seq3: n=66; Ordinary one-way ANOVA).

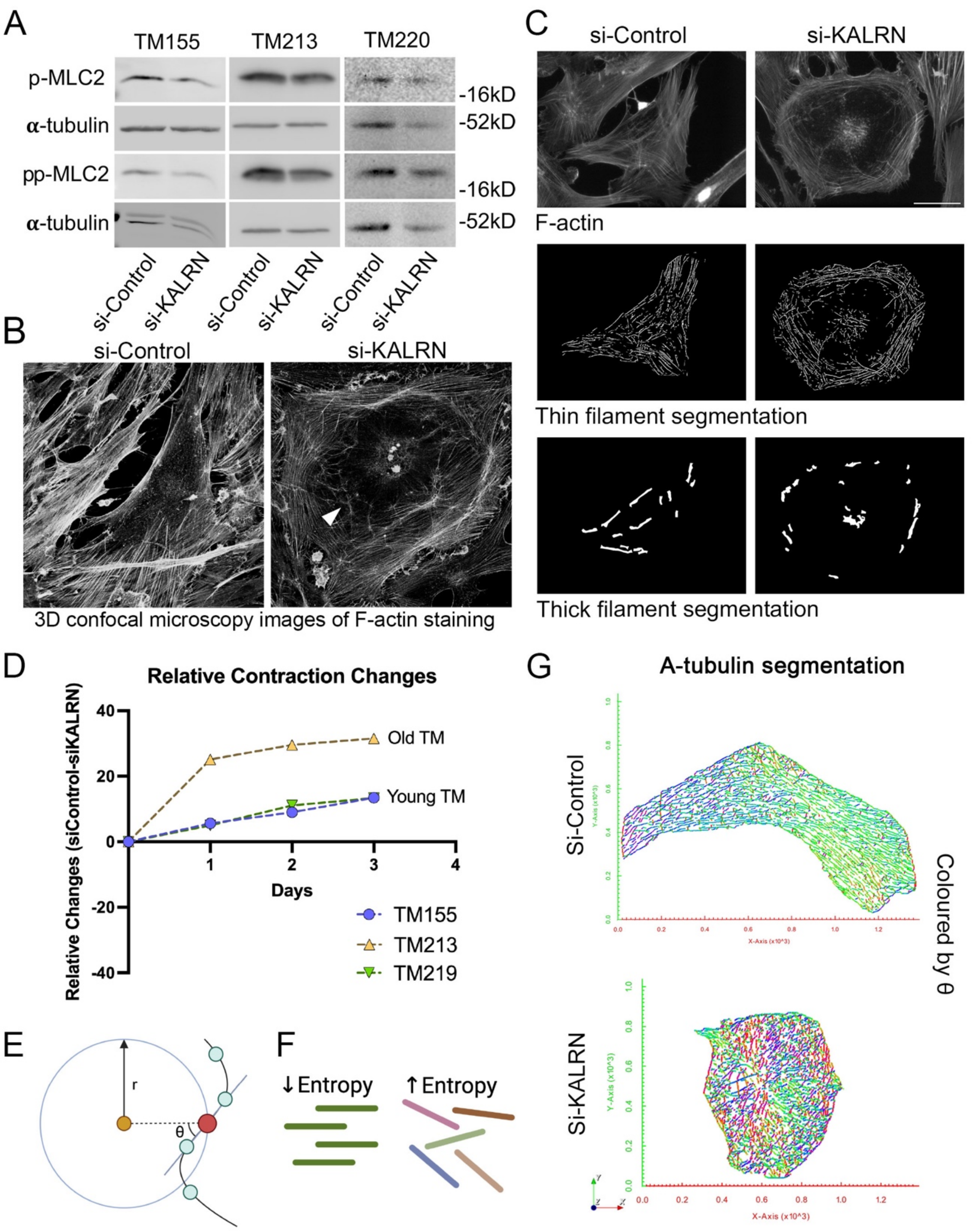

**Supplementary Figure 2: KALRN depletion alters cytoskeletal organization and contractility in TM cells.**

**(A)** Western blotting showing pp-MLC2 and p-MLC2 downregulation in TM cells with KALRN depletion across three additional donors (TM155, TM213, TM220), supporting the quantification in Fig.4B–C. **(B-C)** Immunostaining of F-actin (grey) in KALRN-depleted or control TM cells showing cross-linked actin networks (CLANs; white arrow), with thin (1–2 pixels) and thick (3–15 pixels) filaments segmented using FSegment. Scale bar 50µm**.** **(D)** Contraction assay of TM cells from one older donor (TM213) and two younger donors (TM155, TM219) following KALRN depletion. A greater decrease in contraction was observed in TM213. Each dot represents the difference between the mean of three technical replicates for si-Control and si-KALRN. **(E-F)** Schematic of α-tubulin filament orientation relative to the radial axis (θ) and corresponding entropy, where higher entropy indicates increased filament disorganization. **(G)** Segmented α-tubulin filaments in KALRN-depleted and control cells, coloured by orientation (θ). Similar colours indicate filaments with similar orientations.

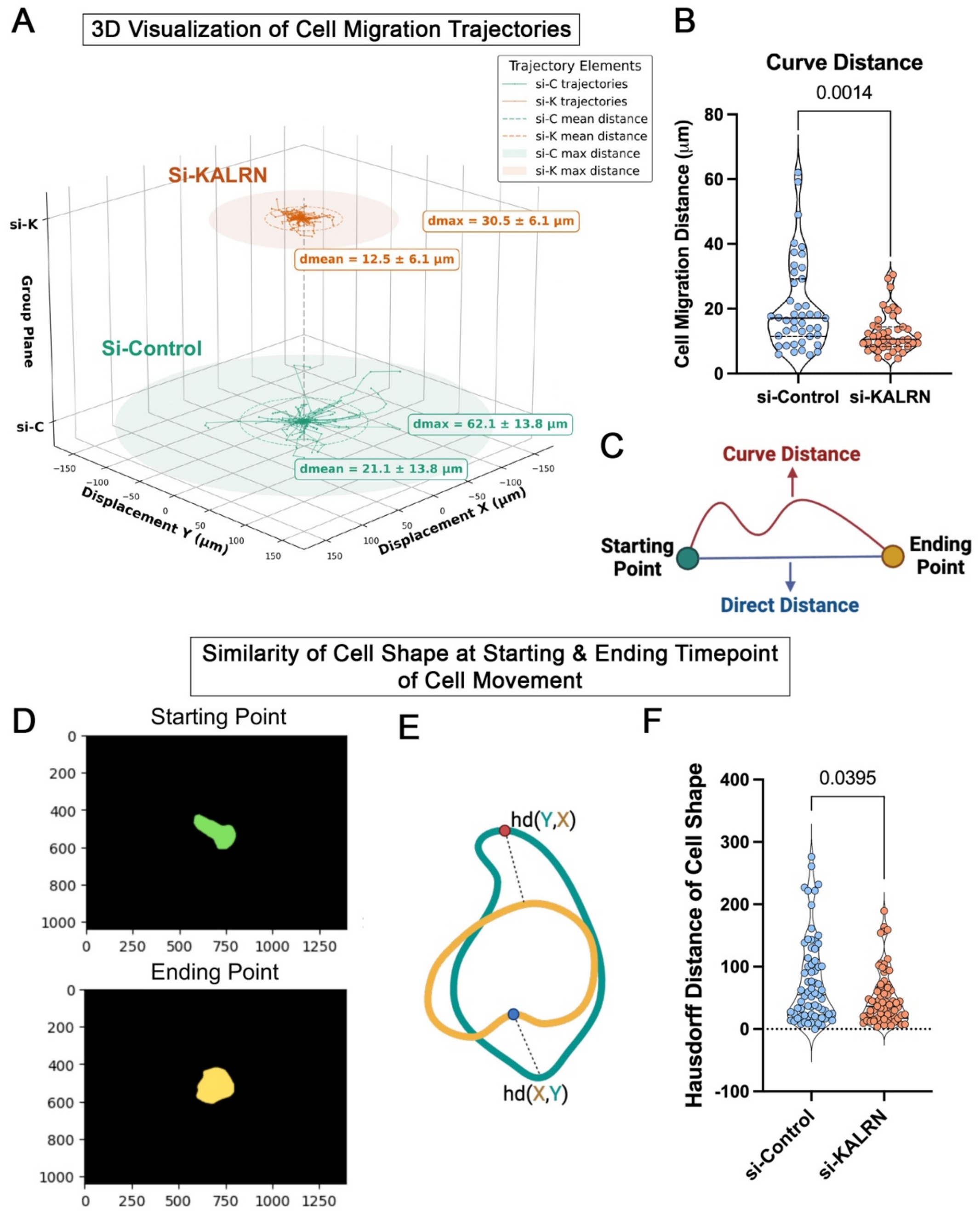

**Supplementary Figure 3: KALRN depletion alters TM cell migration and shape dynamics**

**(A)** 3D visualization of migration trajectories (cell centroids) for KALRN-depleted (orange) and control (green) TM cells from live-cell imaging (Supplementary Videos 1 and 2). Images were acquired every 2 min over 63 frames per field of view, across six fields per group. Trajectories were generated from segmented cell centroids by python packages. Si-KALRN: n=43 cells; Si-Control: n=42 cells. dmean: average straight-line distance from start to end of cell movement; dmax: maximum straight-line distance from start to end. **(B-C)** Curve distance of cell migration from start to end points for KALRN-depleted and control TM cells. KALRN depletion reduced migration distance. Sample sizes: si-KALRN, n=43 cells; si-Control, n=42 cells. Mann–Whitney test. **(D-F)** Similarity of cell shape for single cells from starting to ending points by quantifying the Hausdorff distance (HD) **D,** Central alignment of the segmented cells at starting and ending timepoints for HD calculation. **E,** Schematic of Hausdorff distance (HD) calculation. HD quantifies the maximum dissimilarity between cell outlines at two time points, with lower values indicating greater shape similarity. X represents the coordinates of the cell outline at the starting time point, and Y represents the coordinates at the ending time point. **F,** HD between starting and ending time points was reduced in KALRN-depleted cells, indicating greater cell shape changes in control cells (n=69 cells for each group; Mann–Whitney test).

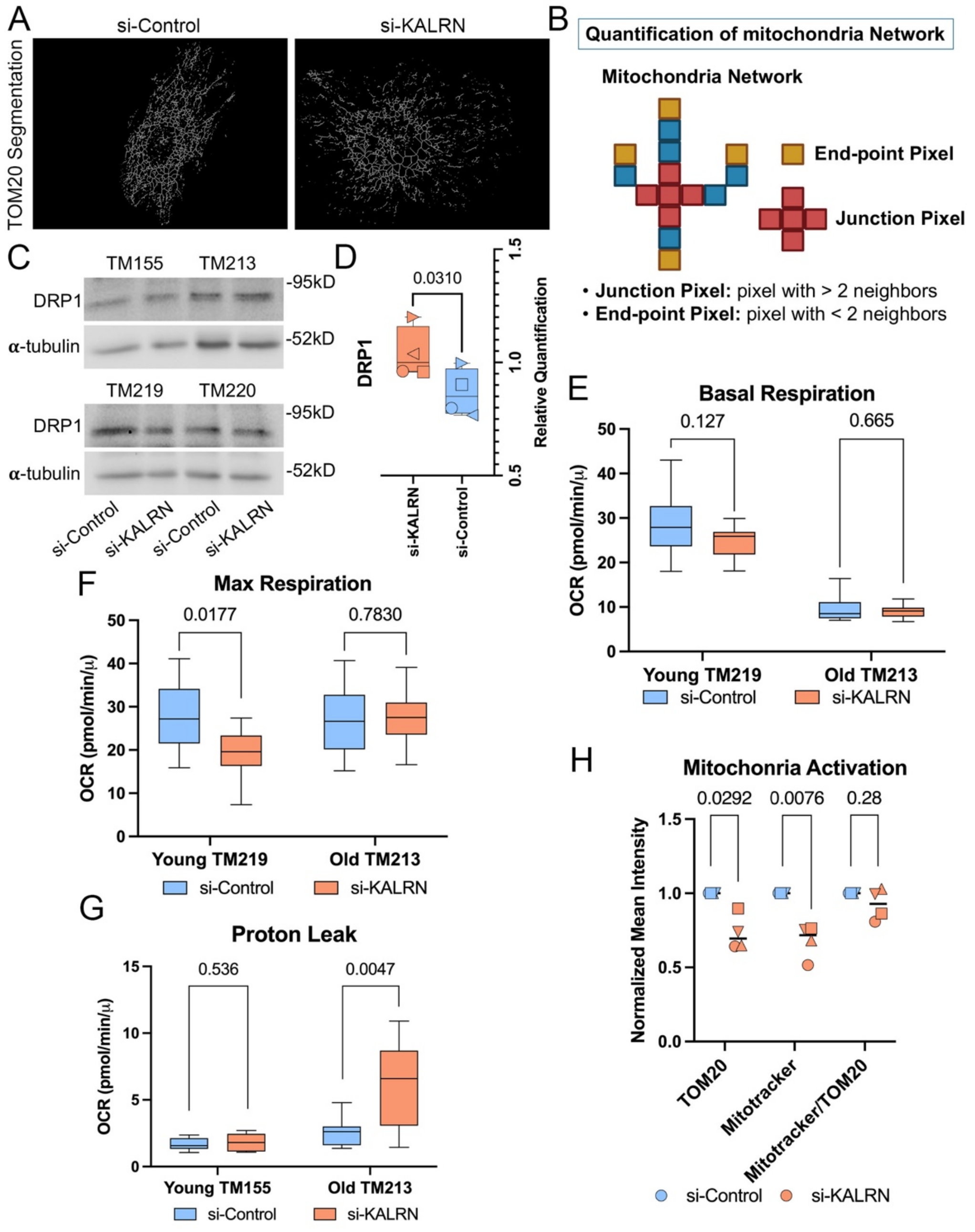

**Supplementary Figure 4: KALRN depletion disrupts mitochondrial network organization and alters mitochondrial activity.**

**(A)** Segmentation of TOM20-positive mitochondrial networks in control and KALRN-depleted cells using ImageJ/Fiji, showing increased fragmentation in KALRN-depleted cells. **(B)** Diagram of junction and end-point pixel detection used to measure mitochondrial network integrity with the AnalyzeSkeleton plugin in ImageJ/Fiji. **(C-D)** Western blotting showing increase DRP1 protein (78-82kDa) expression in KALRN depleted TM cells indicating activated mitochondrial fission (n=4 donors; paired t-test, two tailed). **(E)** Basal mitochondrial respiration was not significantly changed in KALRN-deficient cells from a young donor (TM219; n=12 technical replicates; unpaired t-test) or an old donor (TM213; n=11 technical replicates; Mann–Whitney test). **(F)** Maximum mitochondrial respiration was reduced in KALRN-deficient cells from a young donor (TM219; n=12 technical replicates; Mann–Whitney test) and was not significantly changed in cells from an old donor (TM213; n=12 technical replicates; unpaired t-test). **(G)** Mitochondria proton leak was increased in KALRN-deficient cells from an old donor (TM213; n=11 technical repeats; unpaired t-test) and showed no significant change in cells from a young donor (TM155; n=9 technical replicates; Mann–Whitney test). **(H)** Mitochondrial activity assay in KALRN-depleted cells. Cells were stained with TOM20 and MitoTracker. TOM20 and MitoTracker intensities were reduced in KALRN-depleted groups (n=4 donors; paired t-test).

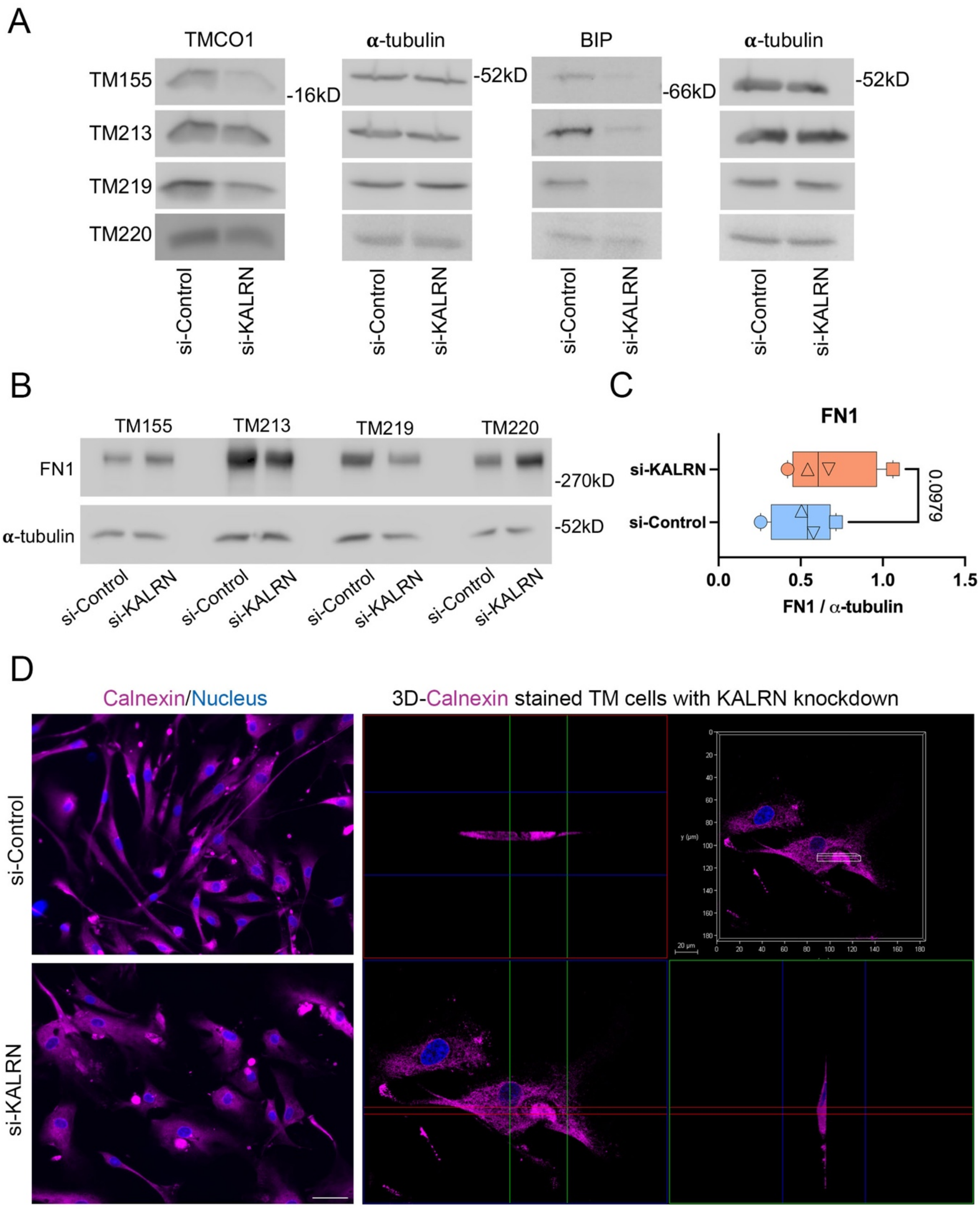

**Supplementary Figure 5: Effects of KALRN depletion on ER and ECM marker expression in TM cells.**

**(A)** Western blotting showing TMCO1 and BIP downregulation in TM cells with KALRN depletion across all four donors, supporting the quantification in Fig.3B–C. **(B-C)** Western blotting and quantification showing an increase of FN1 in TM cells with KALRN depletion across all four donors. Paired t-test. **(D)** Immunofluorescence showing Calnexin (purple) in vacuole-like expansions of TM cells with KALRN depletion. Left: scale bar 50μm; right: 3D reconstruction, scale bar 20 μm.

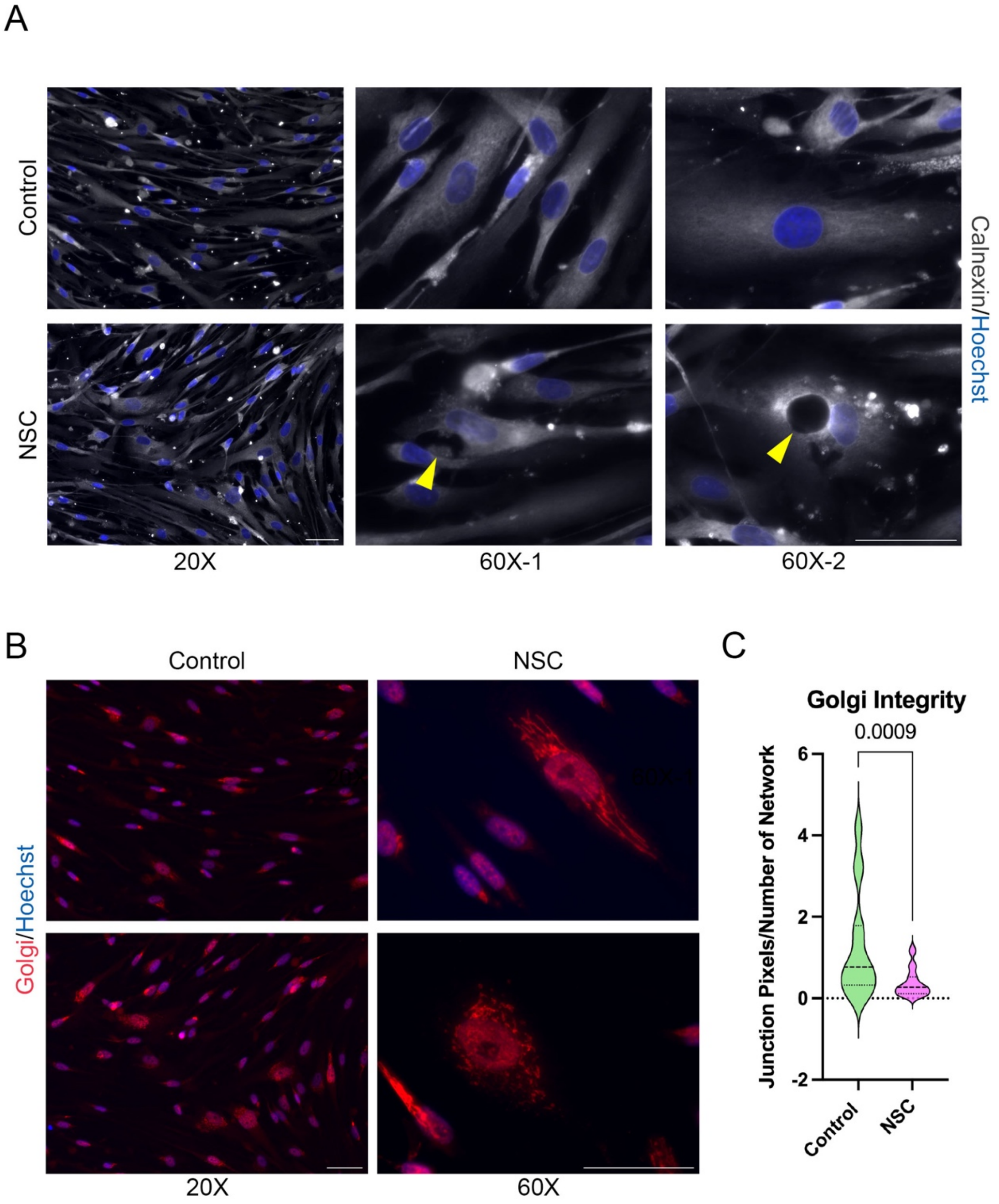

**Supplementary Figure 6: RAC inhibition causes vacuole accumulation and Golgi structure disruption in TM cells resembling KALRN depletion.**

**(A)** Immunostaining of ER marker Calnexin showing the large vacuole formation (yellow arrows) in the human TM cells with RAC inhibitor NSC**.** Scale bar 50µm. **(B-C)** Immunostaining of Golgi marker GOLPH2/GP73 and quantification showing the decreased Golgi structure integrity in KALRN depleted TM cells (TM219); Sample sizes: control: n=28 cells; NSC: n=38 cells. Mann–Whitney test. Scale bar 50µm.

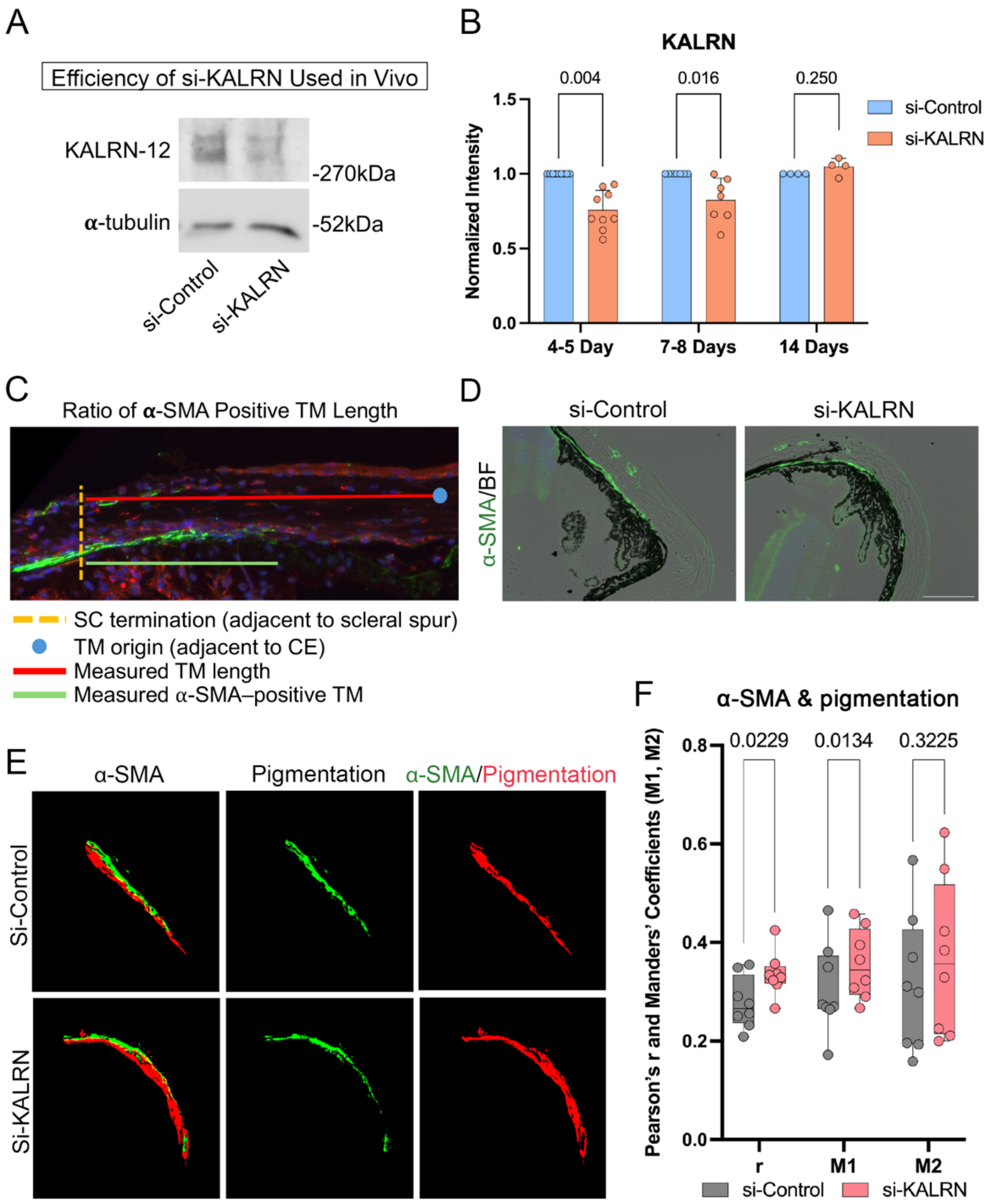

**Supplementary Figure 7: KALRN depletion reduces KALRN expression in mouse TM and enhances α-SMA expression in pigmented TM regions**

**(A)** Western blotting showing efficient depletion of KALRN-12 (~340kDa) in cultured mouse kidney cells using Accell KALRN siRNA, the same reagent applied *in vivo*. **(B)** Quantification of KALRN staining in mouse TM tissue following siRNA injection (4–5 days; n=9 mice; 7–8 days, n=7 mice; day 14, n=4 mice; Wilcoxon test). **(C)** Diagram illustrating TM regions analysed for α-SMA–positive length (see Fig.8H–I). TM was defined between the end of the corneal endothelium (TM origin) and Schlemm's canal (SC) termination near the scleral spur. SC location was identified by α-SMA co-staining with TM/SC markers. α-SMA–positive TM (green) was measured from SC termination to the end of positive staining toward the TM origin. **(D-F)** Co-localization of α-SMA and pigmentation in TM with or without KALRN depletion. **D**, Merged images of α-SMA staining (green) and brightfield channel (pigmentation, brown) to assess overlap. Scale bar, 100µm. **E,** Segmentation of α-SMA positive and pigmented TM regions. **F,** Co-localization quantified with Pearson's correlation (r) and Manders' coefficients (M1, M2). KALRN depletion increased α-SMA/pigmentation overlap (r=1, perfect; r=0, none). Results suggest that pigmented TM regions contained more α-SMA (increased M1), while the proportion of pigmented tissue overlapping α-SMA remained unchanged (M2). Interpretations: M1=1, all α-SMA overlaps with pigmentation; M1=0, no overlap; M2=1, all pigmentation overlaps with α-SMA; M2=0, no overlap. N=8 mice; paired t-test.

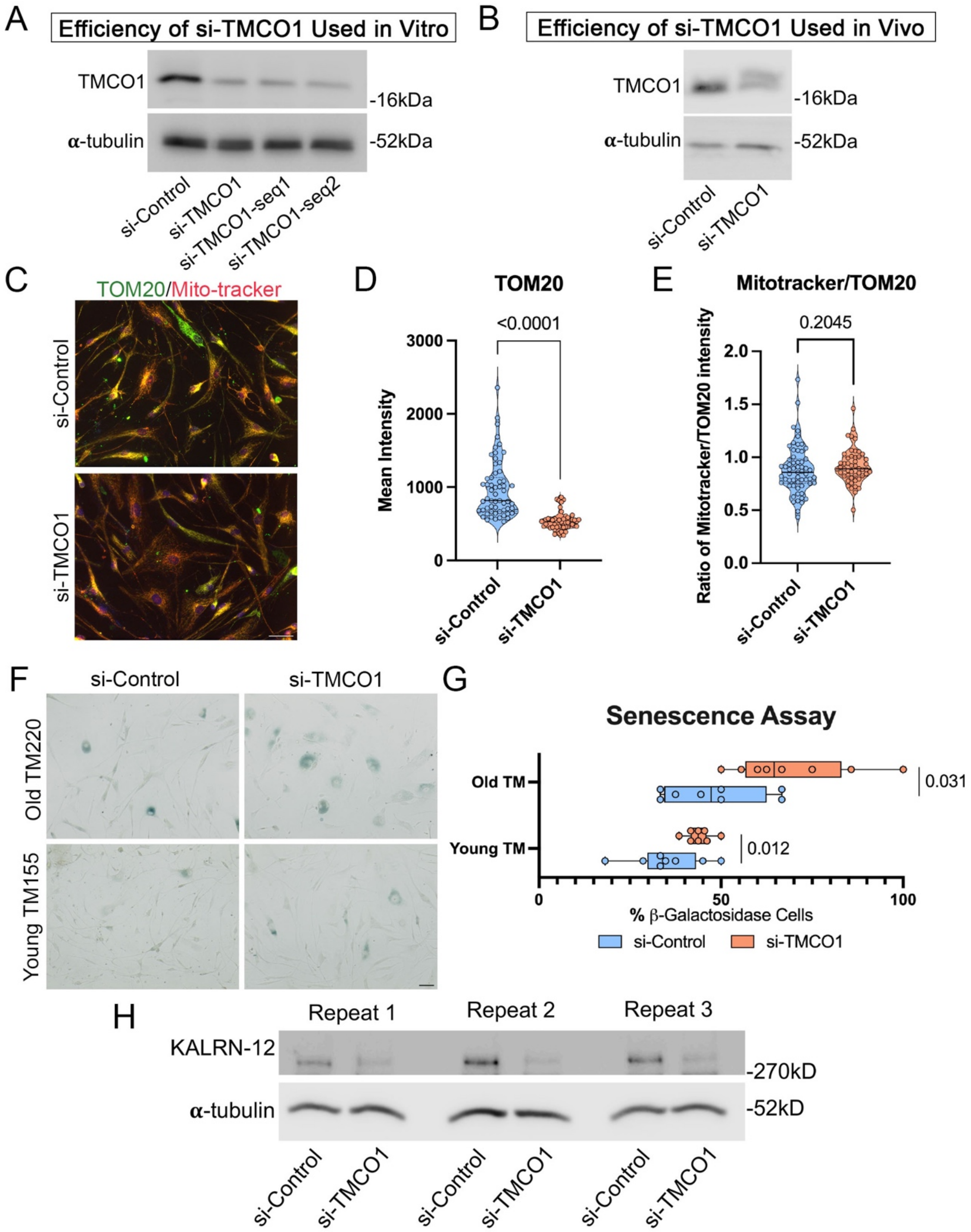

**Supplementary Figure 8: TMCO1 depletion disrupts mitochondrial markers, promotes senescence in TM cells, and reduces KALRN-12 expression**

**(A-B)** Western blotting showing efficient depletion of TMCO1 in cultured human TM cells (TM155) and mouse kidney cells using TMCO1 siRNAs or Accell TMCO1 siRNA, the same reagent applied *in vitro* and *in vivo*. **(C-E)** Immunostaining of TOM20 (green) and MitoTracker (red) in human TM cells after TMCO1 depletion. **C:** TMCO1 depleted TM cells stained with TOM20 (green) and Mito-tracker (red). Scale bar, 50μm. **D:** Quantification of TOM20 staining intensity showing reduction in TMCO1-depleted cells (si-Control, n=59; si-TMCO1, n=49; Mann-Whitney test). **(E)** MitoTracker/TOM20 ratio, reflecting mitochondrial activity, remained unchanged (Mann-Whitney test). **(F-G)** β-Galactosidase senescence assay showing increased proportion of senescent cells (blue) in TMCO1-depleted TM cells. Analysis included n=8 images per group, repeated in two donors; unpaired (t-test). Scale bar, 50μm. This supplements Fig.9H–I, including an additional donor and donor comparisons. **(H)** Western blotting showing decreased KALRN-12 (~340kDa) expression in TMCO1-depleted cells (three technical repeats as in Fig.9K).

| Donor number | Age | Sex | Race |
|---|---|---|---|
| TM155 | 58 | F | White |
| TM213 | 81 | M | White |
| TM219 | 21 | M | White |
| TM220 | 74 | F | White |

| Antibody name | Company | Catelogue | Species | Dilution-ICC | Dilution-WB |
|---|---|---|---|---|---|
| A-catenin | Sigma | C2081 | Rabbit | 1:1000 | 1:2000 |
| ZO-1 | Life technologies | 339100 | Mouse | 1:1500 | 1:1000 |
| I-Afadin | Invitrogen | PA1-25075 | Rabbit | 1:500 | 1:2000 |
| N-cadherin | Invitrogen | 33-3900 | Mouse | 1:100 | N/A |
| BIP | Cell signaling | 3177s | Rabbit | 1:100 | 1:1000 |
| Calnexin | Abcam | ab219644 | Goat | 1:100 | N/A |
| Climp | N/A | N/A | Mouse | 1:100 | N/A |
| KALRN | Invitrogen | PA5-36953 | Rabbit | 1:100-300 | 1:500 |
| TMCO1 | Abcam | ab220729 | Rabbit | 1:100 | 1:1000 |
| Ki67 | Invitrogen | 7B11 | Mouse | 1:100 | N/A |
| P-myosin | Cell signaling | 3675L | Mouse | 1:400 | N/A |
| Pp-myosin | Cell signaling | 95777S | Rabbit | 1:400 | 1:1000 |
| A-SMA | Sigma | A2547 | Mouse | 1:500 | 1:1000 |
| A-tubulin | Abcam | ab18251 | Rabbit | 1:2000 | N/A |
| Tom20 | Santa cruz | SC-17764 | Mouse | 1:100 | 1:500 |
| LAMP1 | Abcam | ab25630 | Mouse | 1:100 | 1:1000 |
| LC3 | Cell signaling | 4108S | Rabbit | 1:100 | 1:1000 |
| C-myc | Santa cruz | F2719 | Mouse | 1:200 | N/A |
| C-myc | MBL | 562-5 | Rabbit | 1:500 | N/A |
| GOLPH2/GP73 (luminal domain) | Matter and Balda (affinity purified fusion protein antibody, luminal domain | | Rabbit | 1:200 | N/A |
| IRDye 680LT Donkey anti-Rabbit | LI-COR | 926-68023 | Donkey | N/A | 1:10,000 |
| IRDye 800CW Donkey anti-Mouse | LI-COR | 926-32212 | Donkey | N/A | 1:10,000 |
| Donkey anti-Mouse IgG (H+L) Highly Cross-Adsorbed Secondary Antibody, Alexa Fluor™ 488 | Invitrogen | A32766 | Donkey | 1:600 | N/A |
| Donkey anti-Rabbit IgG (H+L) Highly Cross-Adsorbed Secondary Antibody, Alexa Fluor™ Plus 555 | Invitrogen | A32794 | Donkey | 1:600 | N/A |
| Donkey anti-Goat IgG (H+L) Highly Cross-Adsorbed Secondary Antibody, Alexa Fluor™ Plus 647 | Invitrogen | A32849 | Donkey | 1:600 | N/A |
| HRP Donkey anti-Rabbit IgG (dissolved as recommended and then diluted by 50% with glycerol) | Invitrogen | A16035 | Donkey | N/A | 1:5,000 |
| HRP Donkey anti-Mouse IgG (dissolved as recommended and then diluted by 50% with glycerol) | Invitrogen | A16017 | Donkey | N/A | 1:5,000 |
